# Self-Actuating 4D Cell-Strand Bioprinting

**DOI:** 10.64898/2026.07.27.740975

**Authors:** Aixiang Ding, Ana F. Cunha, Mariana B. Oliveira, João F. Mano, Eben Alsberg

## Abstract

Engineering biomimetic tissues with dynamically evolving 3D architectures represents an important direction for next-generation tissue engineering, as it enables recapitulation of the continuous morphogenesis of native tissues during development and regeneration. Here, a self-actuating 4D cell-strand bioprinting platform is developed to engineer complex tissue architectures through autonomous cell contractile force (CCF)-driven morphing without requiring external stimuli. The platform integrates a mechanically compliant and self-softening base hydrogel with embedded high-density cell strands printed using a fast-degrading carrier bioink. During culture, the carrier bioink rapidly degrades while the encapsulated cells proliferate and establish connected cellular networks, generating localized contraction that drives programmable shape transformation. Through spatial patterning of embedded cell strands, constructs with diverse morphologies, including V-shaped, helical, folded, and tubular architectures, are generated via controllable self-actuated morphogenesis. The platform further enables engineering of cartilage-like and bone-like tissues with well-defined curvature configurations and mechanically robust tissue matrices. In addition, programmable multi-tissue engineering is demonstrated through fabrication of a muscle–tendon junction-mimicking construct containing spatially organized fibroblast and myoblast compartments. This self-actuating 4D bioprinting strategy enables highly programmable and directionally controlled morphogenesis using a simple construct design with low cell amount requirements, providing a versatile platform for engineering dynamic tissue architectures.

## 1. Introduction

Biological tissues possess highly complex and hierarchical three-dimensional (3D) architectures that are essential for physiological functions such as mass transport, cell communication, and mechanical support^[1]^. Complex tissue curvature patterns arise during morphogenesis through the coordinated interplay of cellular mechanics, cell-cell and cell-matrix interactions, biochemical signaling, and extracellular matrix (ECM) remodeling^[2]^. Such processes generate spatially heterogeneous mechanical stresses and instabilities, including apical constriction-driven bending, differential growth, and growth-induced buckling, which collectively direct the formation of complex 3D tissue architectures during development^[3]^. Therefore, reproducing the dynamic curvature formation of native tissues in a controllable morphogenetic manner is not only important for understanding fundamental developmental biology, but also essential for engineering functional tissue models that better recapitulate the physiological microenvironment and regenerative capacity of native tissues^[4]^. These demands have stimulated growing interest in next-generation four-dimensional (4D) tissue engineering strategies^[5]^.

4D bioprinting, which integrates smart biomaterials with additive manufacturing technologies, has emerged as a promising approach for engineering living cell-laden constructs capable of programmed shape transformation over time^[6]^. Among various biomaterials, hydrogels are particularly attractive for 4D bioprinting because of their excellent cytocompatibility, ECM-mimicking characteristics, and stimuli-responsive behaviors^[7]^. In most current 4D hydrogel systems, stress mismatches are encoded within them through strategies such as multilayer material assembly, spatial material patterning, or functional gradient incorporation^[8]^. As a result, printed constructs can undergo programmable shape changes in response to external stimuli such as temperature, light, pH, chemical triggers, electric fields, or magnetic fields^[9]^. Although these approaches have demonstrated effectiveness in inducing shape morphing, external stimulus-driven systems often require specialized stimulation devices, which may limit their accessibility and translational potential. More importantly, these external stimuli can suffer from limited cytocompatibility, insufficient spatiotemporal control in complex biological environments, and potential interference with cell functions, tissue maturation, and long-term in vivo applications.

In contrast, cell-mediated self-actuation utilizes intrinsic cellular contractility to drive autonomous and biologically adaptive shape transformations that more closely mimic native tissue morphogenesis and long-term tissue remodeling^[10]^. Developing cell-based actuation strategies in 4D bioprinting, particularly through harnessing cell contractile forces (CCFs), is therefore highly attractive because it enables biomimetic and physiologically relevant tissue morphogenesis without relying on external stimulation^[10c^, ^10d^, ^11]^. Compared with externally triggered systems, cell-mediated actuation provides intrinsically cytocompatible, programmable, and adaptive mechanical forces that facilitate the formation of complex tissue architectures, functional maturation, and regenerative tissue remodeling.

Despite its significant potential, enabling effective CCF-based 4D bioprinting remains highly challenging because the contractile forces generated by cells must overcome the mechanical resistance of the printed construct matrix^[12]^. Individual cellular contractile forces are relatively weak, typically on the order of tens of nanonewtons^[13]^, whereas bioprinted constructs generally require sufficient mechanical robustness to maintain post-printing structural stability and printing fidelity. Balancing these competing requirements between mechanical integrity and cell-driven deformability therefore represents a major challenge in the field. To address this issue, two primary strategies have been explored^[10c^, ^10d^, ^11^, ^14]^. The first strategy involves designing mechanically compliant or self-softening hydrogels using degradable polymer networks or sacrificial components, thereby maintaining initial structural stability while permitting gradual cell-induced deformation during culture^[10c, 10d, 14]^. The second strategy focuses on enhancing cell-cell and cell-matrix interactions by incorporating cell-adhesive ligands into hydrogel networks and increasing cell density (e.g., up to 100 × 10^6^ cells/mL bioinks) within constructs to promote collective cellular contraction^[10c, 10d, 11a, 14b, 14c]^. In some cases, these two approaches are combined synergistically to amplify cell-induced actuation^[10c, 10d]^. In addition, these systems must simultaneously support long-term cell survival, proliferation, differentiation, and ECM deposition while preserving sufficient structural fidelity throughout progressive tissue morphogenesis. Moreover, achieving spatially programmable morphogenesis typically requires complex multilayer constructs with spatially defined contractility differences, substantially increasing fabrication complexity^[10c, 11b]^. Consequently, achieving precise and programmable shape transformation through CCF-mediated actuation continues to be a major challenge despite its enormous promise for engineering biomimetic tissue architectures.

Herein, we report a self-actuating 4D cell-strand bioprinting platform for engineering complex tissue architectures through programmed shape transformation driven by localized CCF-induced contraction (**Scheme 1**). The platform involves printing a mechanically compliant, self-softening, and slow-degrading hydrogel matrix as the base construct, followed by embedded printing of high-density cell strands using a fast-degrading carrier bioink. Upon photo-crosslinking and subsequent culture, strong localized CCFs generated within the embedded cell strands induce controlled morphogenesis, resulting in constructs with well-defined curvature configurations. Furthermore, when stem cells are cultured under tissue-specific differentiation conditions, engineered tissues with complex architectures and functional maturation can be generated. Unlike existing CCF-based systems, which typically require high cell densities and/or complex construct designs, this platform enables programmable morphogenesis using substantially reduced cells while maintaining a simple fabrication strategy. Its high spatial controllability allows complex architectures to be generated from single-layer constructs, an ability that has been difficult to achieve with previously reported CCF-driven systems. This self-actuating 4D bioprinting strategy provides a promising platform for morphogenetic tissue engineering and may advance the development of biomimetic tissue constructs for regenerative medicine applications.

**Scheme 1.**
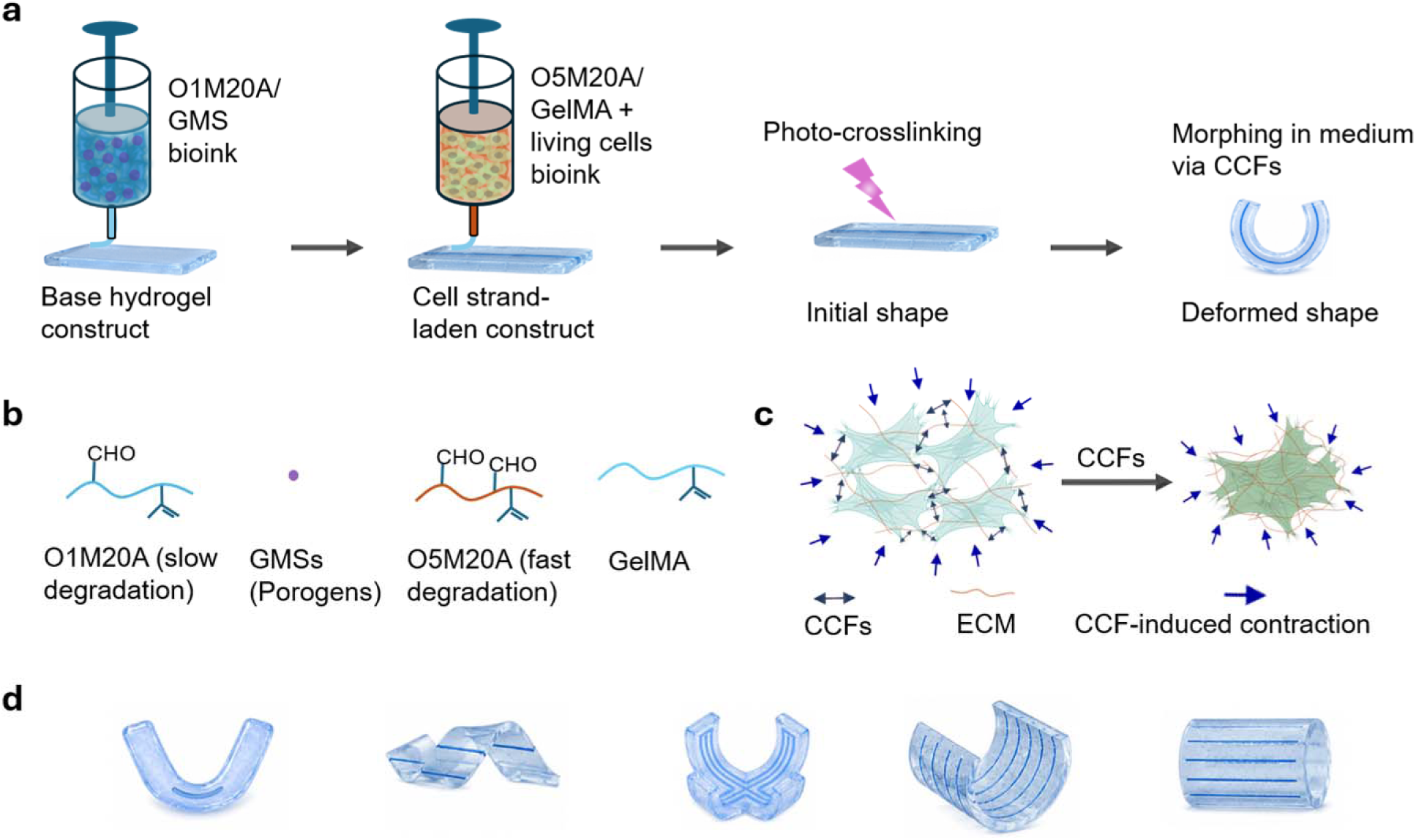
Self-actuating 4D cell-strand bioprinting. (a) Schematic illustration of cell-strand bioprinting and subsequent shape morphing of the printed constructs. (b) Main components used for bioink formulation. (c) Schematic illustration of CCF-mediated contraction. (d) Representative examples of complex engineered architectures.

## 2. Results and Discussion

### 2.1. Engineering Differentially Degradable Bioinks for Self-Actuating 4D Bioprinting

To enable self-actuating shape morphing through embedded cell strands, two bioinks with distinct degradation profiles were developed (**Scheme 1a, b**). Oxidized and methacrylated alginate (OMA), possessing tunable degradability and photocrosslinkability^[15]^, was synthesized as the primary polymer for both bioinks. Since the degradation rate of OMA positively correlates with its oxidation level, OMA with theoretical 1% oxidation and 20% methacrylation (termed O1M20A) was synthesized to formulate a slow-degrading bioink for printing the base hydrogel construct, whereas OMA with theoretical 5% oxidation and 20% methacrylation (termed O5M20A) was synthesized to formulate a fast-degrading bioink for high-density cell encapsulation and embedded cell-strand printing. Both OMAs were processed into jammed microgel assemblies using our previously established calcium ion (Ca^2+^) crosslinking–mechanical fragmentation–high-speed centrifugation protocol^[16]^.

To generate a mechanically compliant and self-softening matrix for the base hydrogel construct, gelatin microspheres (GMSs) were incorporated into the O1M20A bioink to form a composite O1M20A/GMS bioink. The GMSs function as sacrificial porogens because they can liquefy during culture at 37 °C, which has previously been shown to soften the hydrogel matrix from several kilopascals to several hundred pascals^[10c, 17]^. In addition, to enhance initial cell-matrix interactions within the fast-degrading bioink, GelMA containing cell-adhesive RGD ligands was mixed with O5M20A to form O5M20A/GelMA carrier bioinks. The fast-degradation design could enable initial extrusion printing stability while subsequently permitting efficient cell proliferation, migration, and organization during matrix degradation.

Rheological characterization of O1M20A/GMS, O5M20A/GelMA, and cell-laden O5M20A/GelMA (O5M20A/GelMA/cells) bioinks was performed to evaluate their suitability for extrusion-based printing, which requires bioinks to exhibit shear-thinning behavior through a solid-to-fluid phase transition under applied shear stress, followed by rapid self-healing to restore their solid-like state after extrusion to maintain post-printing structural stability. Frequency sweep tests conducted at 0.1% strain showed that the storage modulus (G′) remained higher than the loss modulus (G″) over the frequency range of 0.1–10 Hz for all three bioinks, indicating predominantly solid-like behavior (**Figure S1**). The shear-thinning properties of the bioinks were evaluated by monitoring modulus changes under increasing shear strain and viscosity changes under increasing shear rates. O1M20A/GMS (**Figure 1a**) and O5M20A/GelMA (**Figure 1b**) exhibited crossover points between G′ and G″ at strains of 16% and 79%, respectively, together with decreasing viscosity at increasing shear rates (**Figure 1d, e**), confirming their shear-thinning behavior and suitability for extrusion through printing needles.

**Figure 1.**
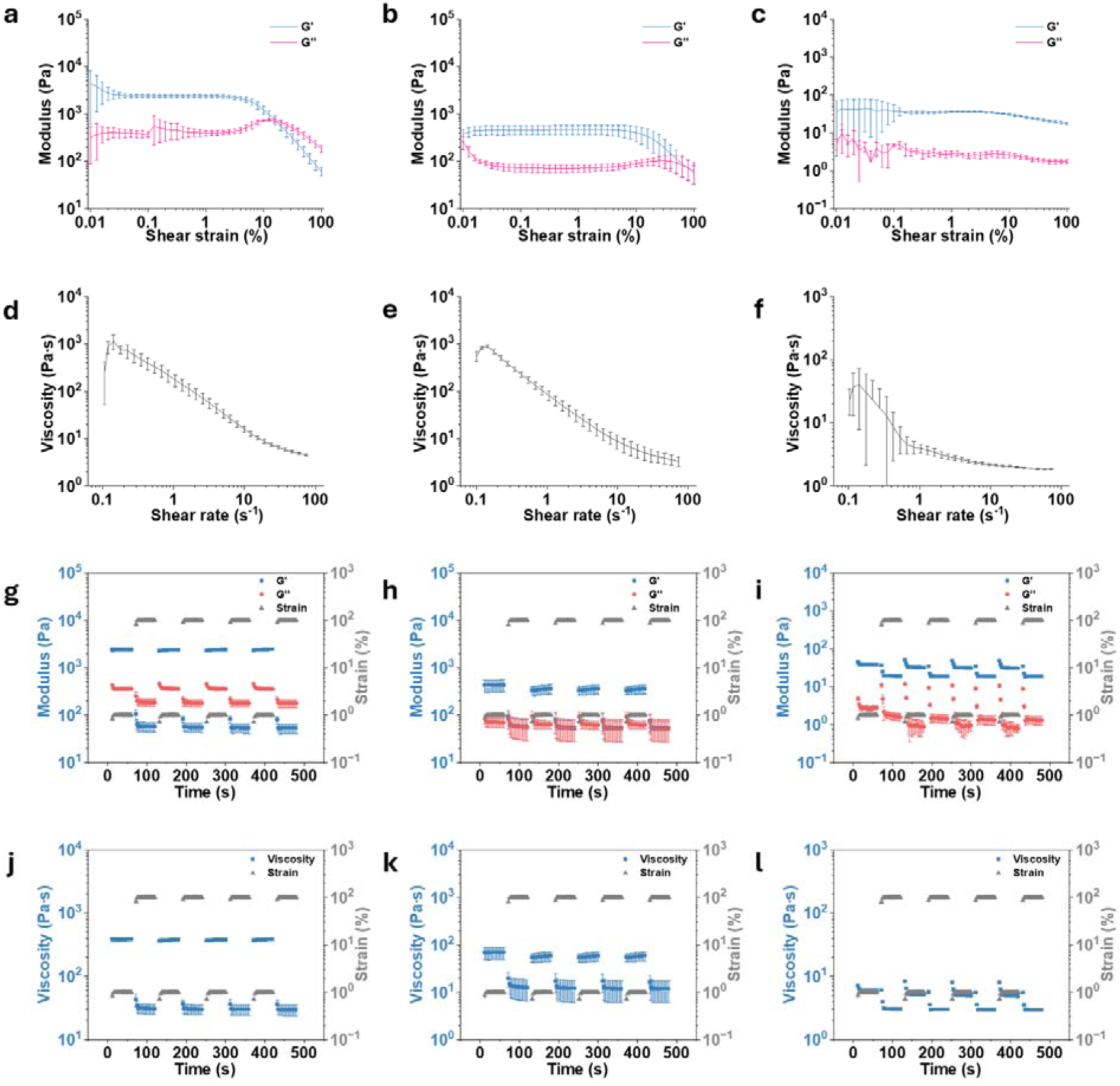
Rheological characterization of bioinks. (a-c) Storage modulus (G′) and loss modulus (G″) as functions of shear strain. (d-f) Viscosity as a function of shear rate. (g-i) G′ and G″ and (j-l) viscosity under cyclic strain switching between 1% and 100%. (a,d, g, j) O1M20A/GMS bioinks; (b, e, h, k) O5M20A/GelMA bioinks; and (c, f, I, l) O5M20A/GelMA/cells bioinks. NIH3T3 cells were encapsulated at a density of 200 million cells mL^-1^ bioink. Data are presented as mean ± standard deviation (±SD), *N* = 3.

When NIH3T3 cells were encapsulated within O5M20A/GelMA at a high density of 200 million cells mL^-1^ bioink, the resulting O5M20A/GelMA/cells bioink exhibited consistently higher G′ than G″ without a clear modulus crossover (**Figure 1c**). Nevertheless, the bioink displayed fluid-like characteristics with an extremely low G′ range of 17–44 Pa. In addition, viscosity decreased with increasing shear rate (**Figure 1f**), likely because cells deform and align along the flow direction under shear stress, thereby reducing flow resistance, a phenomenon commonly observed in concentrated cell suspensions^[18]^. This fluid-like behavior facilitates smooth extrusion, while the O5M20A/GelMA matrix provides sufficient mechanical support to maintain homogeneous cell suspension and prevent sedimentation, enabling the formation of cell strands with evenly distributed cells.

The self-healing properties of the bioinks were evaluated by cyclic strain switching between 1% and 100% strain while monitoring changes in modulus and viscosity. O1M20A/GMS and O5M20A/GelMA exhibited rapid and reversible switching between solid-like (G′ > G″ at 1% strain) and fluid-like (G′ < G″ at 100% strain) states (**Figure 1g, h**), accompanied by corresponding reversible viscosity transitions (**Figure 1j, k**), without noticeable signal fatigue during repeated cycling. These results demonstrate their excellent self-healing capability^[19]^. Although O5M20A/GelMA/cells did not exhibit a crossover between G′ and G″ because of the predominance of cells within the bioink, cyclic reductions in both G′ and G″ under 100% strain were observed (**Figure 1i**), together with reversible viscosity changes during strain cycling (**Figure 1l**), indicating recoverable rheological behavior and self-healing capability. The excellent self-healing behavior of O1M20A/GMS enables rapid recovery after extrusion to form stable freestanding constructs, whereas the self-healing behavior of O5M20A/GelMA/cells facilitates recovery to a colloidally stable state after embedded printing within the O1M20A/GMS matrix.

Collectively, the rheological properties of O1M20A/GMS, including shear-thinning and rapid self-healing, support its suitability for extrusion printing of the base hydrogel construct, while the fluid-like behavior and recoverable rheological properties of O5M20A/GelMA/cells make it highly suitable for embedded cell-strand printing.

To investigate the role of GMSs in softening the base hydrogel construct, photocrosslinked O1M20A/GMS composite hydrogels were mechanically characterized in the as-prepared state and after swelling at room temperature (RT) and 37 °C (**Figure 2a, b**). Compared with the as-prepared hydrogels, which exhibited a G′ of 2.75 kPa at 1 Hz, swollen hydrogels showed a substantial decrease in G′. Swelling at 37 °C resulted in a further significant reduction in G′ to approximately 480 Pa, likely due to dissolution and removal of GMSs that weakened the hydrogel network, consistent with our previous reports on GMS-incorporated hydrogels^[10c]^. The mechanical properties of the O1M20A/GMS hydrogels could also be tuned by varying the photocrosslinking duration. Increasing the UV crosslinking time from 10 s to 30 s gradually increased G′ (**Figure 2c, d**), providing a useful strategy for modulating morphability, including morphing kinetics and deformation extent (discussed later).

**Figure 2.**
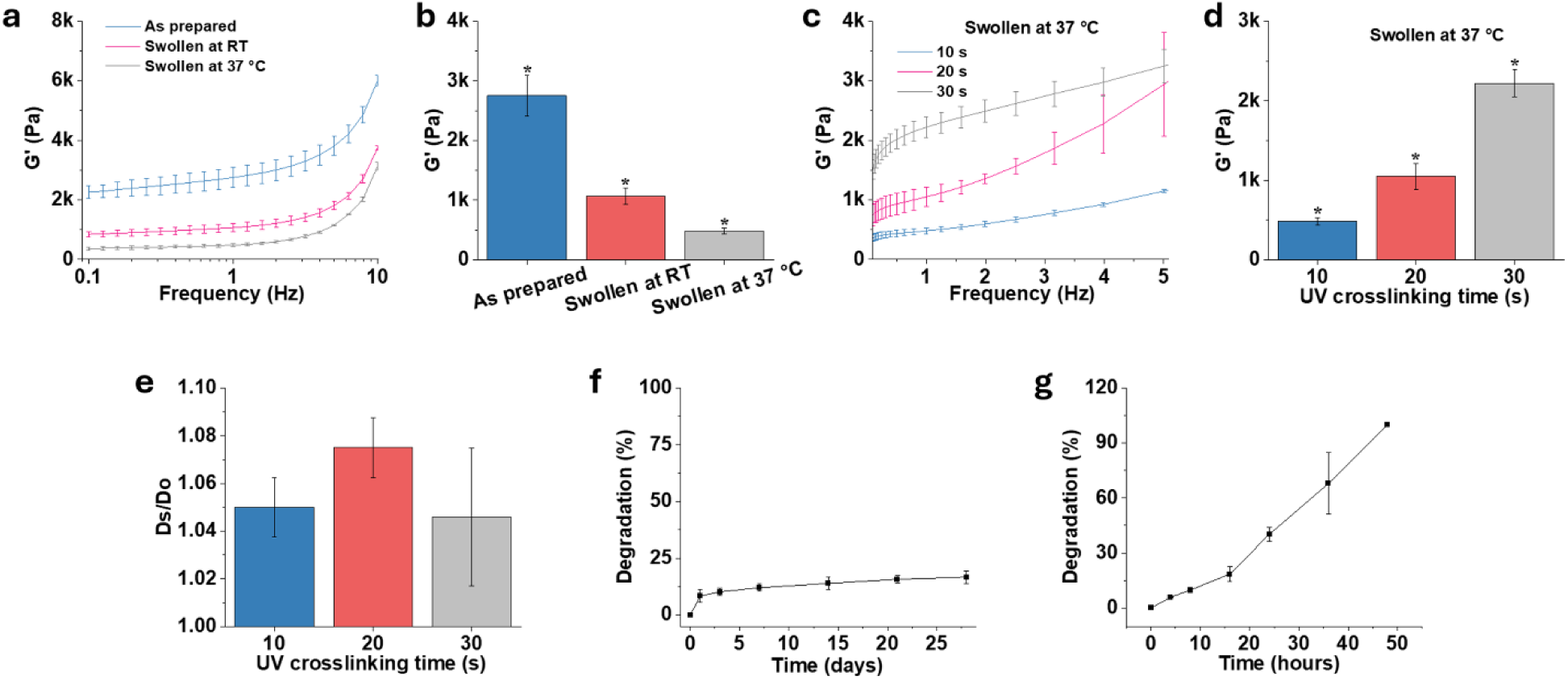
Mechanical, swelling, and degradation characterization of crosslinked hydrogels. (a) G′ of O1M20A/GMS hydrogels as a function of oscillatory frequency and (b) G′ of O1M20A/GMS hydrogels at 1 Hz under different culture conditions; hydrogels were prepared with a 10 s UV crosslinking time and cultured in medium for 12 h prior to testing. (c) G′ of O1M20A/GMS hydrogels as a function of oscillatory frequency and (d) G′ of O1M20A/GMS hydrogels at 1 Hz after 10–30 s UV crosslinking and subsequent swelling at 37 °C. (e) Swelling behavior of O1M20A/GMS hydrogels crosslinked for 10–30 s. Ds denotes the swollen hydrogel diameter, while Do denotes the original hydrogel diameter. Degradation of (f) O1M20A hydrogels (10 s UV crosslinking) and (g) O5M20A/GelMA (10 s UV crosslinking) over culture time. Data are presented as mean ± standard deviation (±SD), *N* = 3.

Despite differences in photocrosslinking duration, all O1M20A/GMS hydrogels exhibited minimal swelling, with a D_s_/D_o_ ratio close to 1.0 and no significant differences among groups (**Figure 2e**). Degradation studies further revealed distinct degradation profiles between the two hydrogel systems. O1M20A hydrogels remained structurally stable with minimal degradation over 28 days of culture (**Figure 2f**), whereas O5M20A/GelMA hydrogels underwent rapid degradation and were completely degraded within 48 h of culture (**Figure 2g**). The stable O1M20A/GMS matrix therefore provides long-term structural support during tissue culture, while the rapid degradation of O5M20A/GelMA is expected to generate biomaterial-free cell strands with enhanced cell-cell interactions and amplified collective CCFs.

### 2.2. Single Cell-Strand Actuation Enables Programmable Shape Morphing

O5M20A/GelMA/cell bioinks could be successfully incorporated into the O1M20A/GMS base hydrogel through embedding printing, forming uniform cell strands (**Figure S2**). To investigate the role of embedded cell strands in driving construct morphing, strip-shaped hydrogel constructs (14 mm × 2.5 mm × 1.0 mm, length × width × thickness) containing single embedded cell strands were fabricated and cultured in regular growth medium (GM) to monitor shape transformation over time. Cell types including NIH3T3 fibroblasts, C2C12 myoblasts, human mesenchymal stem cells (hMSCs), and human umbilical vein endothelial cells (HUVECs) were evaluated. In addition, the influence of matrix mechanical properties, regulated by varying UV crosslinking duration (10–30 s), was investigated.

For comparison, three control groups were included: an “Empty” group without embedded cell strands, a “CytoD” group treated with the CCF inhibitor cytochalasin D (CytoD)^[20]^, and a “Dead” group containing dead cell strands. Constructs in all control groups exhibited minimal shape changes during 7 days of culture, whereas constructs containing NIH3T3 strands showed progressively increased bending over time (**Figure 3a**). Hydrogels crosslinked with shorter UV exposure times exhibited faster bending kinetics and larger bending angles compared with those crosslinked for longer durations throughout the culture period (**Figure 3b**), eventually forming ring- or “C”-shaped constructs by Day 7 (D7). At D7, the embedded cell strand remained intact and clearly visible within the construct (**Figure S3**), while the formed construct remained soft yet mechanically robust enough to withstand vigorous stirring with a spatula (**Movie S1**).

**Figure 3.**
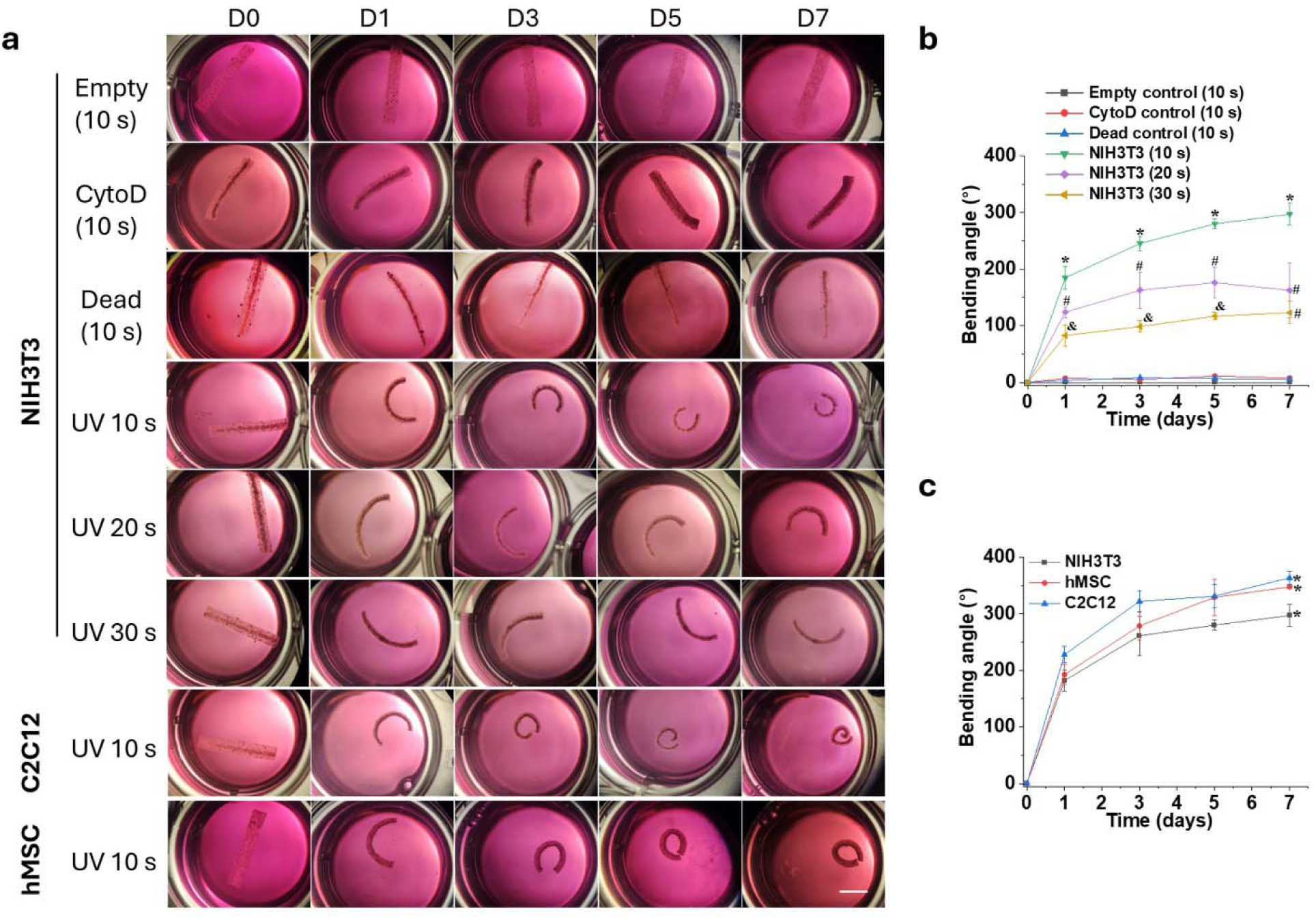
Shape-morphing characterization of hydrogel strips. (a) Representative images showing the progressive bending of hydrogel strips during 7 days of culture in GM. (b) Bending angles of empty control hydrogel strips and NIH3T3 strand-laden hydrogel strips over time. *, ^#^, ^&^*p* < 0.05 compared among groups with different labels and the three control groups (Empty, CytoD, and Dead). (c) Bending angles of NIH3T3-, hMSC-, and C2C12 strand-laden hydrogel strips over time. \**p* < 0.05 at D7 between the NIH3T3 group and the other two groups. Scale bar = 5 mm. Data are presented as mean ± standard deviation (±SD), *N* = 3.

Among the cell types investigated, constructs containing C2C12 and hMSC strands exhibited significantly larger bending angles at D7 compared with constructs containing NIH3T3 strands (**Figure 3c**). In contrast, constructs containing HUVEC strands exhibited negligible shape morphing during culture (**Figure S4**). These results suggest that the self-actuating morphing behavior strongly depends on cell type-specific contractile activity and the ability of embedded cells to generate sufficient collective contractile forces to overcome matrix mechanical resistance.

### 2.3. Dynamic Cellular Organization Underlies Self-Actuated Shape Morphing

The progressive enhancement of construct bending over time is likely attributed to gradually increased contraction mediated by collective CCFs within the embedded cell strand. To further investigate the role of CCF generation in driving self-actuated morphing, NIH3T3 strand-laden constructs were subjected to live/dead staining and histological analyses at different culture times. Live/dead staining demonstrated predominantly viable cells (green) throughout the culture period (**Figure 4a, S5**). Cells treated with CytoD also remained highly viable, whereas cells in the “Dead” group were completely nonviable (**Figure S6**).

**Figure 4.**
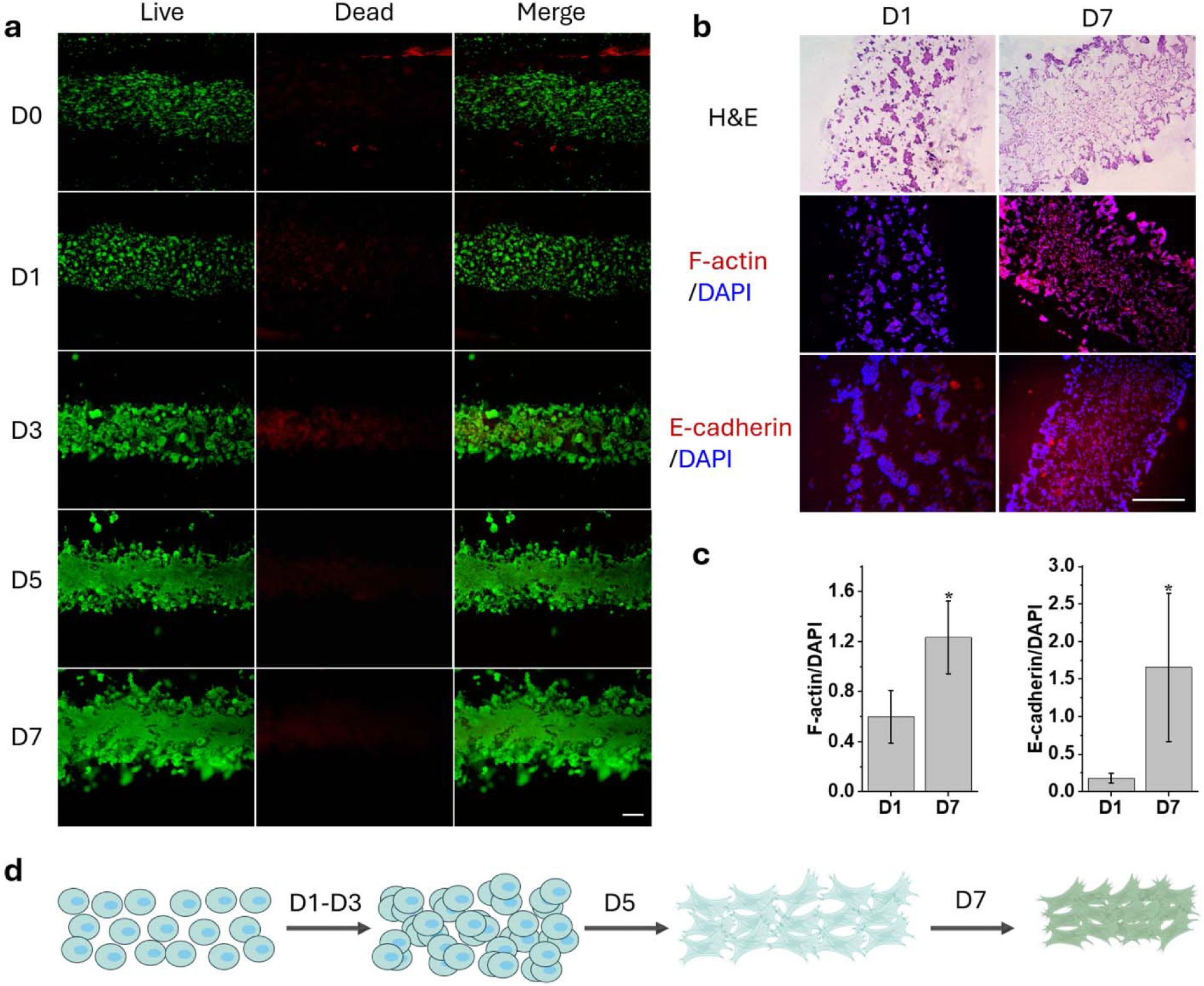
Live/dead and histological characterization of NIH3T3 strand-laden hydrogel strips during culture in GM. (a) Live/dead staining of NIH3T3 strand-laden hydrogel strips. (b) H&E, F-actin, and E-cadherin staining at D1 and D7. (c) Signal intensity ratios of F-actin/DAPI and E-cadherin/DAPI at D1 and D7. (d) Schematic illustration of the temporal changes in cell morphology, cellular distribution, and interconnected network formation within the embedded cell strand. \**p* < 0.05 compared to D1. Scale bars = 0.25 mm. Data are presented as mean ± standard deviation (±SD), *N* = 3.

Interestingly, NIH3T3-laden constructs exhibited a clear temporal transition in cell distribution and morphology, progressing from dispersed individual cells at D0, to cell aggregates at D1 and D3, and ultimately to highly dense and interconnected cellular networks at D5 and D7 (**Figure 4d**). This increase in cellular density was accompanied by an approximately 5-fold increase in DNA content at D7 compared with D0 (**Figure S7**), likely resulting from substantial cell proliferation facilitated by cell-ECM interactions and the rapid degradation of the O5M20A/GelMA carrier matrix. The morphological transition from early-stage cell aggregates to highly organized cellular networks was further confirmed by histological analyses (**Figure 4b, S8**), including hematoxylin and eosin (H&E) staining, phalloidin staining of F-actin, and immunofluorescence staining of E-cadherin. F-actin is a major cytoskeletal component responsible for generating CCFs through actomyosin interactions and transmitting these forces throughout the cellular network and surrounding matrix^[21]^. E-cadherin, a transmembrane glycoprotein critical for cell-cell adhesion, functions as a key component of adherens junctions and is an important mechanosensor and mechanotransducer^[22]^.

Compared with D1, significantly enhanced expression of both F-actin and E-cadherin was observed at D7 (**Figure 4c**), suggesting substantially increased cell-cell interactions and collective CCF generation during culture. In contrast, CytoD-treated cells remained dispersed without forming organized cellular networks and exhibited weak F-actin and E-cadherin staining signals (**Figure S9**). These results suggest that progressive formation of interconnected cellular networks and enhanced collective cell contractility are critical for driving the self-actuated morphogenesis of the constructs.

### 2.4. Spatially Encoded Cell Contractility Directs Complex Shape Formation

With confirmation of the critical role of embedded cell strands in mediating self-actuated morphogenesis, we next explored the fabrication of more complex and programmable architectures through rational strand-patterning design. For example, printing a shortened cell strand within the base hydrogel enabled localized contraction, resulting in folding of the strip into a “V”-shaped construct (**Figure 5a**). In contrast, printing multiple inclined strands (6 strands) within a single strip generated well-defined helical architectures, in which the chirality could be programmed into either right-handed or left-handed helices by controlling the strand hatch angle (**Figure 5b, c**). Furthermore, by altering the initial geometry of the printed construct from a strip to a cross-shaped configuration, a folded cross-shaped architecture could be generated after culture-induced morphing (**Figure 5d**).

**Figure 5.**
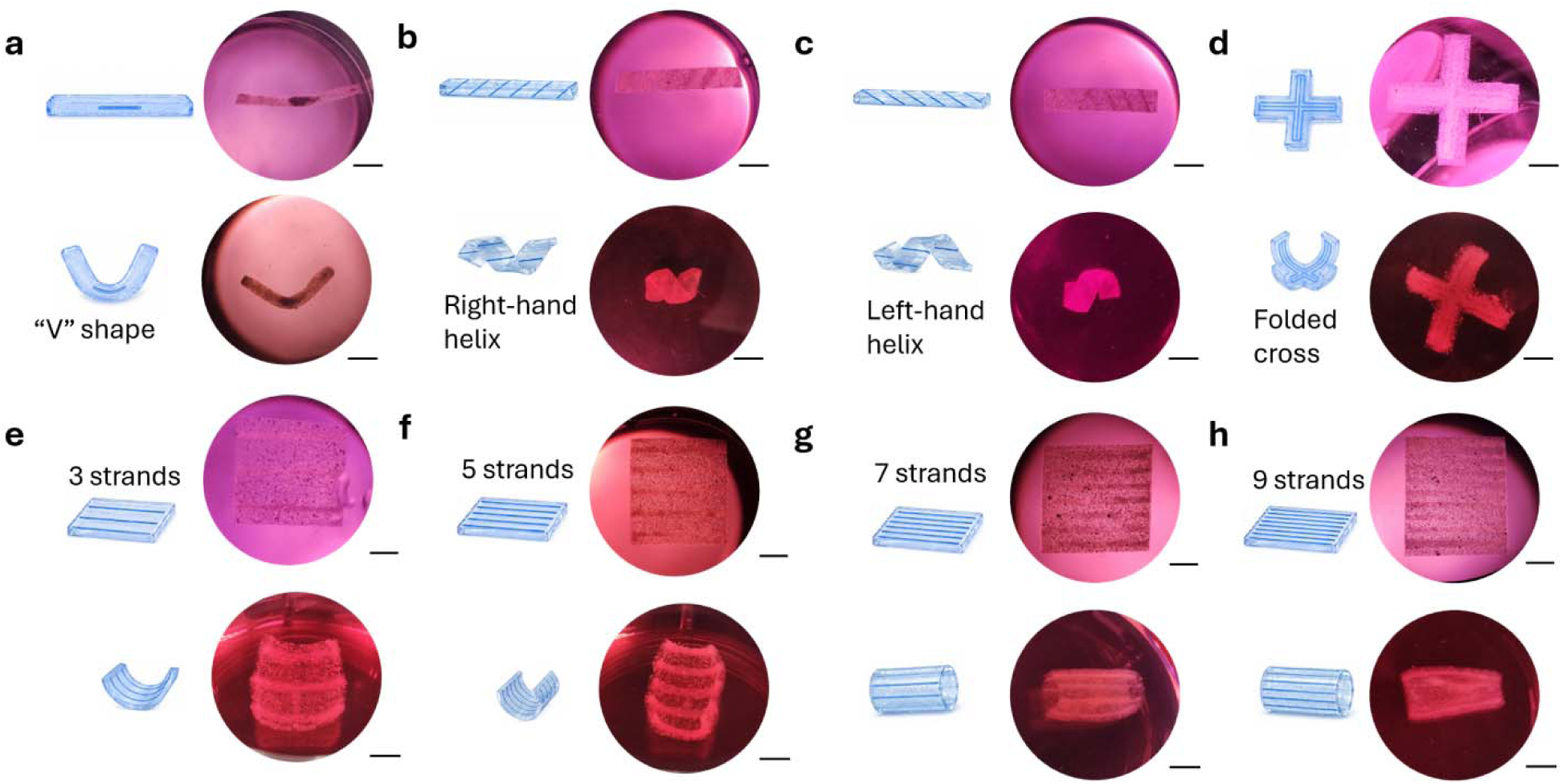
Complex shape engineering of hydrogel constructs. Engineering of (a) V-shaped, (b) right-handed helical, (c) left-handed helical, (d) folded cross-shaped, and sheet-based constructs with (e) 3 strands, (f) 5 strands, (g) 7 strands, and (h) 9 strands. NIH3T3 cells were used, and constructs were cultured for 7 days in GM prior to imaging. Scale bars = 5 mm.

The morphing behavior of larger sheet-like constructs could also be modulated by controlling strand density. Hydrogel sheets containing increasing numbers of embedded cell strands (3–9 strands) were fabricated and cultured to investigate their shape transformation behavior. As the strand number increased from 3 to 5, an enhanced bending curvature along the longitudinal strand direction was observed, likely due to increased overall collective contractility within the constructs (**Figure 5e, f**). Interestingly, when the strand number further increased to 7 or 9, the sheet constructs no longer predominantly bent along the longitudinal direction, but instead underwent rolling along the lateral direction of the strands, resulting in the formation of closed tubular structures (**Figure 5g, h, S10**). This phenomenon suggests that, at higher strand densities, the collective lateral contractile forces generated within the construct may surpass the longitudinal contractility, thereby altering the dominant morphing direction. Although the precise mechanism underlying this transition remains unclear, this intriguing behavior provides a versatile strategy for programming morphing directionality and generating complex 3D architectures through spatial regulation of cell strand organization and collective cellular contractility. These results demonstrate the strong programmability and design flexibility of the self-actuating cell-strand bioprinting platform for engineering complex morphogenetic tissue structures.

### 2.5. Self-Actuating 4D Morphogenetic Tissue Engineering

Developing 4D morphogenetic tissue engineering strategies based on 4D bioprinting is important because native tissues dynamically evolve their architectures during development and regeneration rather than remaining static structures^[23]^. Conventional tissue engineering approaches primarily fabricate predefined geometries but often fail to recapitulate the progressive morphogenesis, curvature formation, and dynamic cell-matrix remodeling observed in living tissues^[24]^. In contrast, 4D bioprinting enables engineered constructs to undergo programmable and time-dependent shape transformation, thereby better mimicking natural developmental processes and tissue maturation^[25]^. Incorporating cell-mediated self-actuation further provides biologically adaptive and intrinsically biomimetic mechanical forces that coordinate tissue morphogenesis, structural organization, and functional matrix deposition. Such self-actuating morphogenetic systems therefore hold strong potential for engineering physiologically relevant tissues with complex architectures and dynamic functional properties for regenerative medicine applications.

### Cartilage-like tissue engineering

To explore the potential of this platform for 4D morphogenetic tissue engineering, proof-of-concept cartilage-like tissue engineering studies were first conducted. hMSC strand-laden constructs were fabricated and cultured in chondrogenic pellet medium (CPM), while counterpart constructs cultured in regular GM served as controls. Shape morphing of strip-shaped constructs was monitored and quantified over a 21-day culture period. Constructs cultured in both GM and CPM underwent progressive dynamic morphogenesis (**Figure 6a**), forming curved tissue constructs with distinct appearances. GM constructs remained relatively translucent, whereas CPM constructs developed a dense white appearance indicative of substantial cartilage-like matrix deposition (**Figure 6b).** Compared with GM constructs, CPM constructs exhibited slower morphing kinetics and smaller bending angles (**Figure 6c**), likely because rapid deposition of cartilage-like ECM increased construct stiffness and mechanically restricted further cell-mediated deformation. Compression testing confirmed significantly greater mechanical strength in CPM constructs than in GM constructs at D21 (33 kPa *vs.* 1.17 kPa; **Figure 6d**). Both GM and CPM constructs remained mechanically stable and resistant to vigorous stirring in medium (**Movies S2 and S3**). Notably, CPM constructs exhibited excellent elasticity and could rapidly recover their original shape after complete compression with tweezers, whereas GM constructs remained softer and stickier and failed to fully recover after compression (**Movies S4 and S5**).

**Figure 6.**
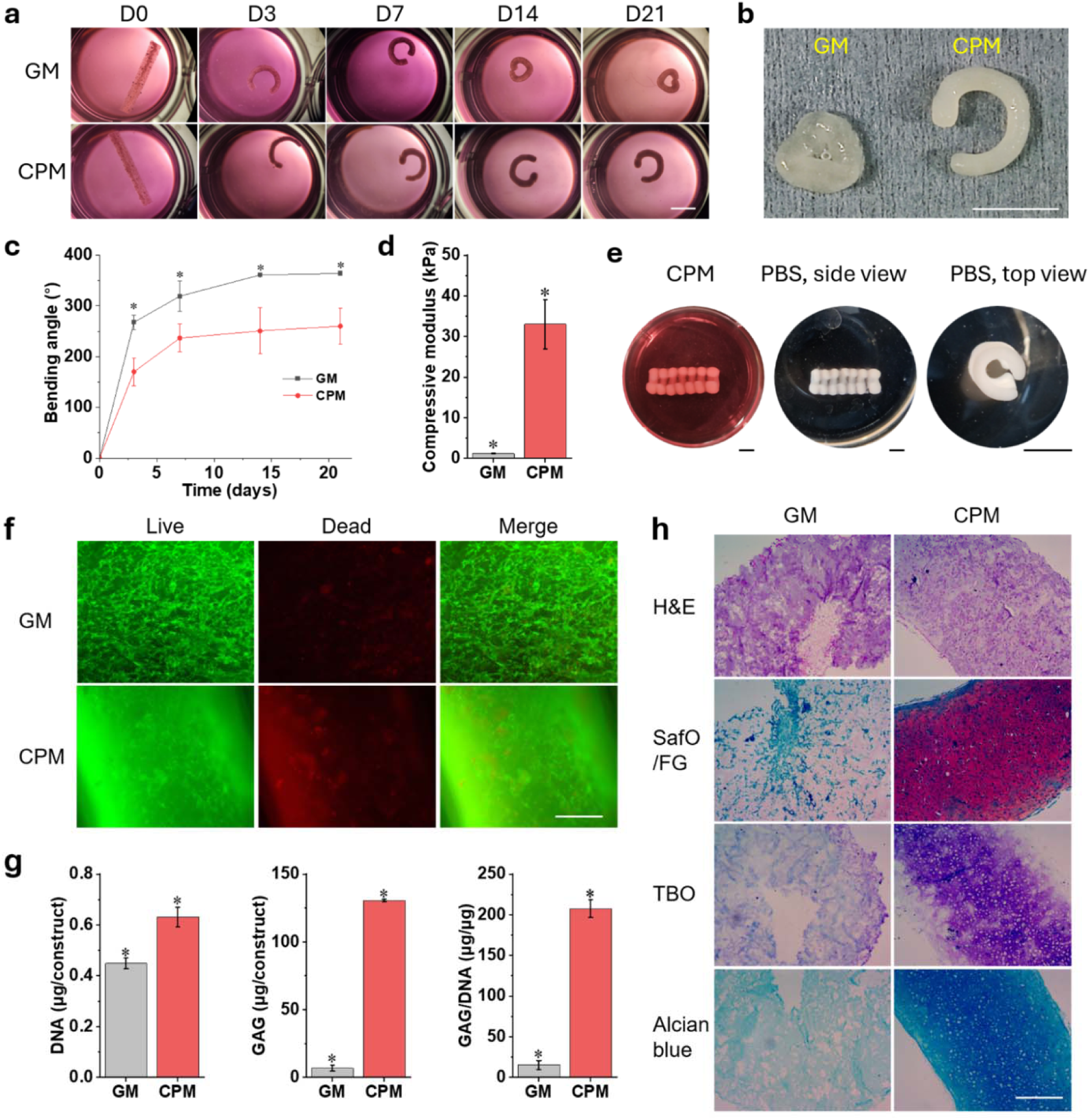
Self-actuating 4D cartilage-like tissue engineering. (a) Representative photomicrographs showing the progressive bending of hMSC strand-laden hydrogel strips during 21 days of culture in GM and CPM. (b) Representative photographs of constructs cultured in GM and CPM at D21. (c) Bending angles of the hydrogel strips cultured in GM and CPM as a function of culture time. (d) Compressive modulus of GM- and CPM-cultured constructs at D21. (e) Photographs of a trachea-mimicking tubular cartilage construct obtained after 21 days of culture in CPM. (f) Photomicrographs of live/dead stain of whole GM and CPM constructs at D21. (g) Biochemical analysis of DNA, GAG, and GAG/DNA content of hydrogel strip constructs at D21. (h) Photomicrographs of H&E, SafO/FG, TBO, and Alcian blue (pH 1.0) stain sections of hydrogel strip constructs at D21. \**p* < 0.05. Scale bars: (a, b, e) = 5 mm; (f, h) = 0.25 mm. Data are presented as mean ± standard deviation (±SD), *N* = 3.

This differentiation strategy could be readily extended to engineer large centimeter-scale biomimetic tissues by incorporating multiple hMSC strands into a single construct prior to differentiation. For example, sheet constructs containing seven embedded strands underwent progressive morphogenesis during 21 days of CPM culture to form trachea-like tubular tissues containing seven “C”-shaped cartilage-like rings within a single construct (**Figure 6e, S11a**). During differentiation, gradual thickening of the cell strands was also observed (**Figure S11b**). The resulting trachea-mimicking tissues remained mechanically robust and resistant to stirring-induced disturbance in medium (**Movie S6**).

Live/dead staining demonstrated high cell viability throughout long-term culture. In GM constructs, highly interconnected cellular networks were clearly visible, whereas cellular staining in CPM constructs was less apparent because the dense cartilage-like matrix hindered dye penetration and imaging (**Figure 6f**). Nevertheless, after prolonged culture, cells had infiltrated the entire strip construct and distinct individual strands were no longer visible. Chondrogenesis was further evaluated through biochemical quantification and histological analyses. DNA content remained significantly elevated in both groups throughout culture, while CPM constructs exhibited significantly higher glycosaminoglycan (GAG) production and GAG/DNA ratios compared with GM constructs (**Figure 6g**), indicating enhanced cartilage matrix formation. Robust cartilage-like tissue regeneration in CPM constructs was further confirmed by intense Safranin O/Fast Green (SafO/FG), toluidine blue O (TBO), and Alcian blue (pH 1.0) staining (**Figure 6h**).

### Bone-like tissue engineering

The applicability of the self-actuating bioprinting platform for 4D morphogenetic tissue engineering was further validated through proof-of-concept bone-like tissue engineering studies. hMSC strand-laden constructs were cultured in osteogenic medium (OM) for 28 days to evaluate shape morphogenesis and osteogenic differentiation, with GM-cultured constructs serving as controls. Similar to the cartilage engineering study, constructs in both groups underwent progressive morphogenesis during culture, forming tissues with distinct appearances in which GM constructs remained relatively translucent whereas OM constructs developed an opaque white appearance (**Figure 7a, b**). OM constructs also exhibited slower morphing kinetics and reduced bending compared with GM constructs (**Figure 7c**), likely because osteoid and mineralized bone-like matrix deposition increased mechanical stiffness and restricted further deformation.

**Figure 7.**
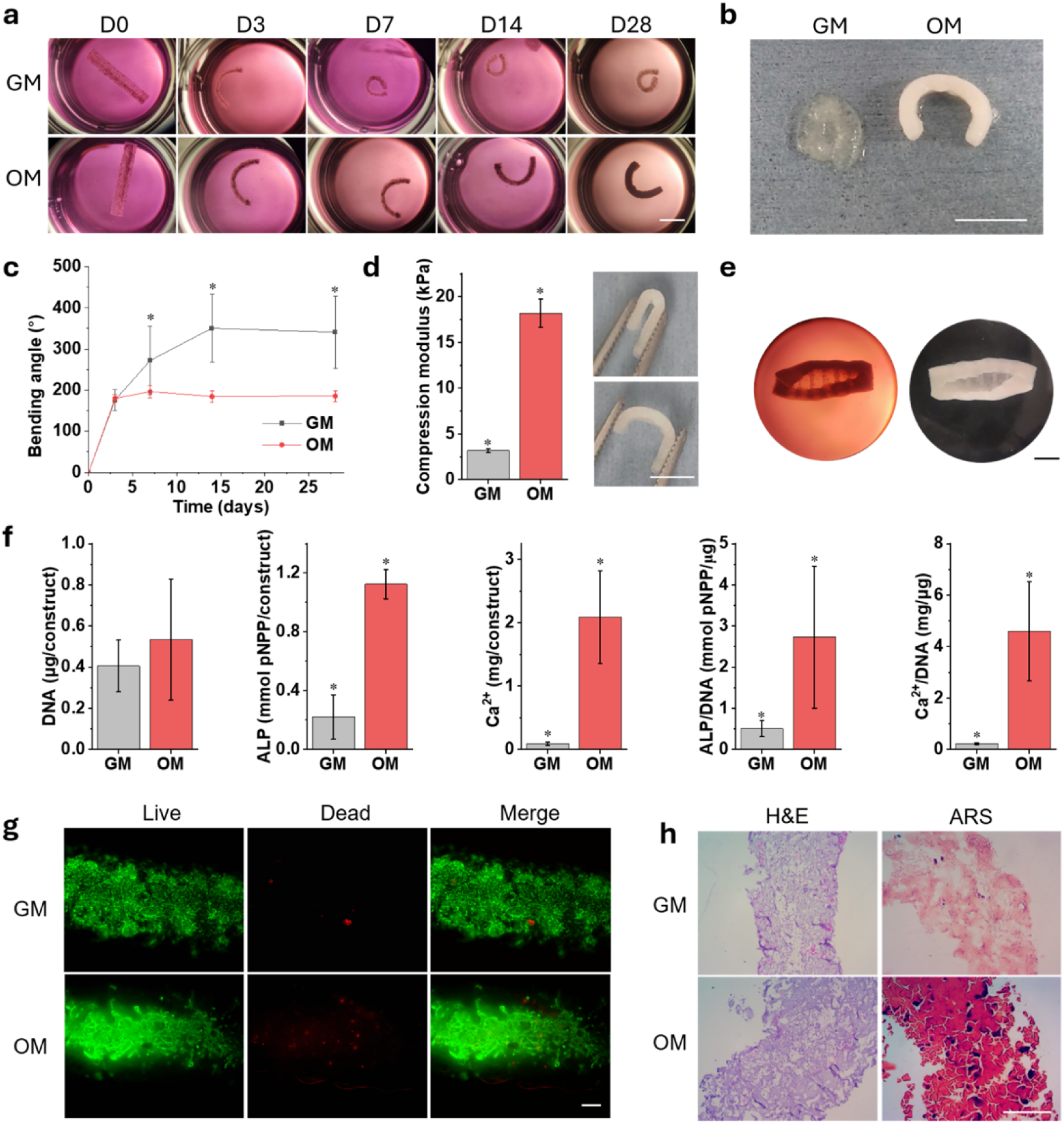
Self-actuating 4D bone-like tissue engineering. (a) Representative photomicrographs showing the progressive bending of hMSC strand-laden hydrogel strips during 28 days of culture in GM and OM. (b) Representative photographs of constructs cultured in GM and OM at D28. (c) Bending angles of hydrogel strips cultured in GM and OM as a function of culture time. (d) Compressive modulus of GM- and OM-cultured constructs at D28. Insets show complete shape recovery of an OM-cultured strip at D28 after full compression with tweezers. (e) Photographs of a tubular bone construct obtained after 28 days of culture in OM. (f) Biochemical analysis of DNA, ALP, Ca^2+^, ALP/DNA, and Ca^2+^/DNA content of hydrogel strip constructs at D28. (g) Live/dead staining of GM and OM constructs at D28. (h) H&E and ARS staining of hydrogel strip constructs at D28. \**p* < 0.05. Scale bars: (a, b, d, e) = 5 mm; (g, h) = 0.25 mm. Data are presented as mean ± standard deviation (±SD), *N* = 3.

Both GM and OM constructs remained mechanically stable during vigorous stirring in medium (**Movies S7 and S8**). However, OM constructs exhibited substantially greater mechanical robustness than GM constructs and could fully recover after complete compression with tweezers, whereas GM constructs could not (**Figure 7d and Movie S9**). Similarly, incorporation of multiple cell strands enabled engineering of larger tissue constructs with well-defined curvature architectures, including mechanically stable tubular constructs containing 7 “C”-shaped bone-like rings (**Figure 7e and Movie S10**). Gradual thickening of the embedded strands during osteogenic differentiation was also observed (**Figure S12**).

Biochemical and histological analyses were further performed to evaluate bone-like tissue formation. Although DNA content in OM constructs was slightly higher than in GM constructs, no significant difference was observed. In contrast, OM constructs exhibited significantly enhanced alkaline phosphatase (ALP) activity, calcium (Ca^2+^) deposition, and corresponding ALP/DNA and Ca^2+^/DNA ratios compared with GM constructs (**Figure 7f**), indicative of effective bone-like matrix formation. Live/dead staining demonstrated high cell viability and dense interconnected cellular networks in both groups at D28 (**Figure 7g**). Successful osteogenic differentiation in OM constructs was further confirmed by strong Alizarin Red S (ARS) staining for mineral deposition (**Figure 7h**).

### Multi-tissue engineering

This self-actuating 4D bioprinting platform also bears strong potential for programmable multi-tissue engineering because the base hydrogel matrix is highly cytocompatible and capable of supporting direct cell encapsulation. To explore this capability, NIH3T3 fibroblasts were encapsulated within the base hydrogel, while embedded C2C12 cell strands were printed to generate a biomimetic myotendinous junction (MTJ)-like construct^[26]^. In this model, C2C12 cells represented the muscle compartment, whereas NIH3T3 fibroblasts served as the tendon-like compartment^[27]^, providing a morphodynamical platform for musculoskeletal tissue engineering.

During 7 days of culture, the NIH3T3-laden base hydrogel alone exhibited minimal shape changes while maintaining high cell viability (**Figure S13**), suggesting that encapsulation of NIH3T3 cells within the base matrix had negligible influence on construct morphing. In contrast, the NIH3T3/C2C12 MTJ-mimicking constructs underwent progressively enhanced bending morphogenesis over time (**Figure 8a, b**). Fluorescence imaging, in which C2C12 cells and NIH3T3 cells were labeled with CellTracker Red and CellTracker Green, respectively, clearly revealed the initial boundary between the embedded C2C12 strand and the surrounding NIH3T3-laden base hydrogel at D0 and D1 (**Figure 8c**). Over time, this boundary gradually became less distinct, likely due to cell proliferation and dilution of fluorescence signals during culture.

**Figure 8.**
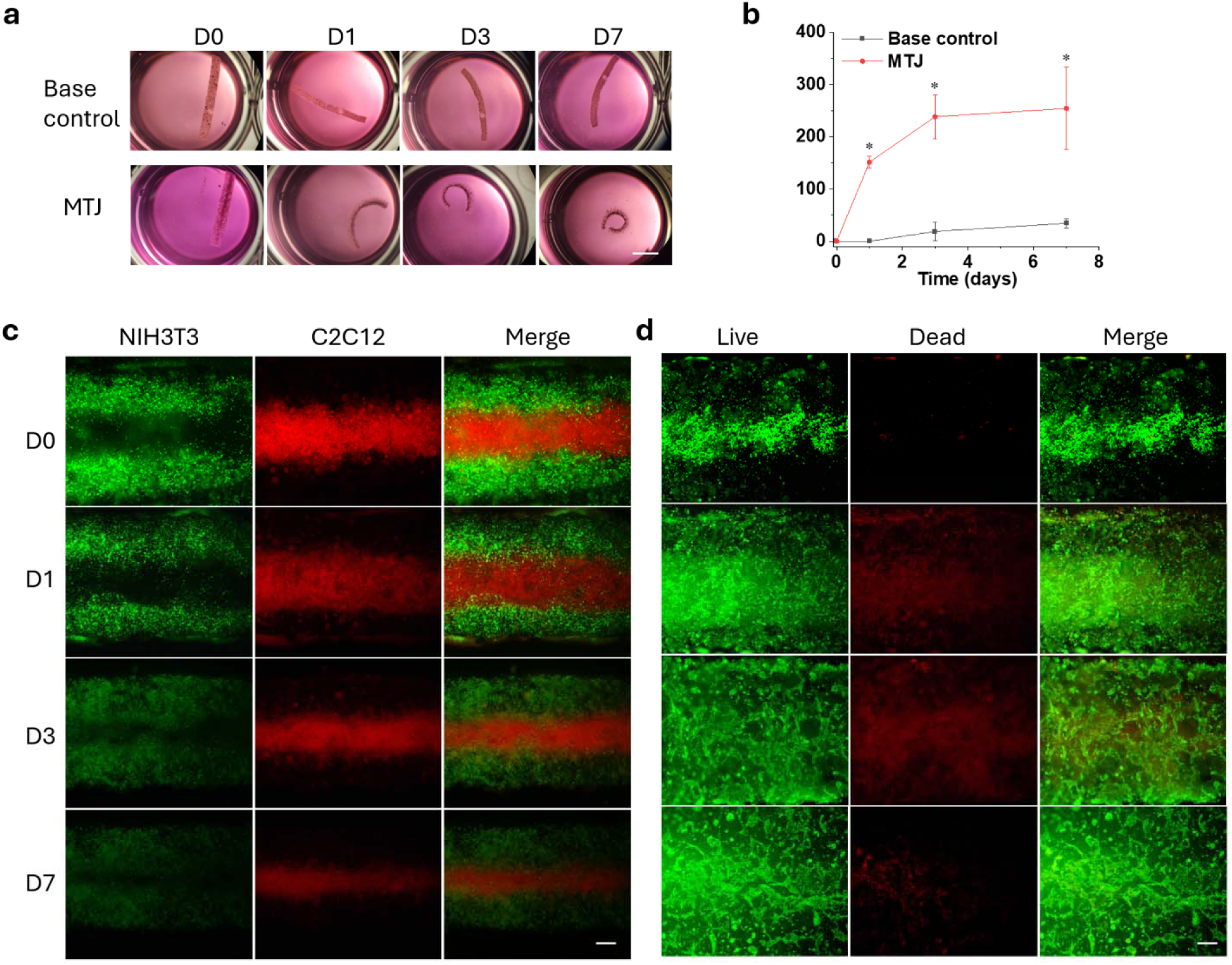
Self-actuating 4D multi-tissue engineering. (a) Representative photomicrographs showing progressive bending morphogenesis of NIH3T3/C2C12 laden hydrogel strips during 7 days of culture in MTJ differentiation medium. (b) Bending angles of the hydrogel strips as a function of culture time. (c) Fluorescence imaging of NIH3T3 (green)/C2C12 (red) laden hydrogel strips at different culture times. (d) Photomicrographs of live/dead stain of NIH3T3 (green)/C2C12 (red) laden hydrogel strips at different culture times. Scale bars: (a) = 5 mm; (c,d) = 0.25 mm. Data are presented as mean ± standard deviation (±SD), *N* = 3.

Importantly, high cell viability was maintained throughout the culture period (**Figure 8d**). In addition, C2C12 cells progressively formed interconnected cellular networks and appeared to extend into the surrounding NIH3T3-laden hydrogel region, which may indicate early-stage MTJ-like tissue integration, potentially resembling myotube extension across the muscle-tendon interface^[28]^. Although further investigation using optimized differentiation conditions, cell densities, and histological characterization will be necessary to fully evaluate MTJ tissue regeneration, this preliminary proof-of-concept study demonstrates that construct morphology can evolve through endogenous CCFs during tissue maturation. Such a dynamic morphogenetic process may provide mechanical cues while simultaneously generating curved or folded muscle–tendon interfaces that resemble a non-flat MTJ architecture. These findings highlight the potential of the self-actuating 4D bioprinting platform for programmable multi-tissue engineering and morphogenetic formation of heterogeneous tissue interfaces.

## 3. Conclusion

In summary, a self-actuating 4D cell-strand bioprinting platform capable of engineering complex tissue architectures through autonomous, CCF-driven morphogenesis without requiring external stimuli was developed. A mechanically compliant and self-softening O1M20A/GMS bioink was engineered for printing the base hydrogel construct, while a fast-degrading O5M20A/GelMA carrier bioink enabled embedded printing of high-density cell strands. The base hydrogel maintained long-term structural stability during culture, whereas rapid degradation of the carrier bioink promoted cell proliferation, cytoskeletal remodeling, and cellular network formation, thereby generating strong collective CCFs that drove progressive shape morphogenesis through localized contraction. Through rational spatial patterning of embedded cell strands, diverse architectures, including V-shaped, helical, folded, and tubular structures, were successfully generated via programmable self-actuated morphogenesis. This strategy achieves highly programmable and directionally controlled morphogenesis using a simple construct design with low cell amount requirements, enabling the generation of complex architectures from single-layer constructs. Importantly, using hMSCs as a differentiation cell source, this platform further enabled 4D morphogenetic tissue engineering of cartilage-like and bone-like tissues with defined curvature configurations, robust ECM deposition, and enhanced mechanical properties. In addition, the cytocompatible nature of the platform enabled programmable multi-tissue engineering, demonstrated through fabrication of a muscle–tendon junction-mimicking construct containing spatially organized NIH3T3 fibroblasts within the base hydrogel and embedded C2C12 myoblast strands. Overall, this study highlights the effectiveness of spatially regulated cellular contractility in driving biomimetic tissue morphogenesis, establishes a versatile biointegrated and self-evolving platform for engineering physiologically relevant tissues with dynamic architectures and functional tissue interfaces, and offers new opportunities for developmental biology studies and organ-scale biofabrication.

## 4. Experimental Section

### Synthesis of O1M20A and O5M20A

O1M20A and O5M20A were synthesized according to previously reported protocols with slight modifications^[15]^. Briefly, alginate (10 g; Protanal LF 120M, apparent viscosity: 251 mPa·S at 1% w/v in water at 25°C, generously provided by Roquette) was dissolved in deionized water (DI H_2_O, 900 mL) under stirring overnight at RT. Sodium periodate (NaIO_4_; 0.108 g for O1M20A and 0.54 g for O5M20A; Sigma-Aldrich) was dissolved in DI H_2_O (100 mL) and rapidly added to the alginate solution. The reaction was maintained under stirring at RT in the dark by covering the container with aluminum foil. Subsequently, 2-(N-morpholino)ethanesulfonic acid (MES; 19.52 g; Sigma-Aldrich) and sodium choloride (NaCl; 17.53 g; Sigma-Aldrich) were added, and the solution pH was adjusted to 6.5 using 5 N sodium hydroxide (NaOH; Sigma-Aldrich). N-hydroxysuccinimide (NHS; 1.18 g; Sigma-Aldrich) and 1-ethyl-3-(3-dimethylaminopropyl) carbodiimide hydrochloride (EDC; 3.89 g; Sigma-Aldrich) were sequentially added, and the reaction proceeded for 10 min. Aminoethyl methacrylate hydrochloride (AEMA; 1.69 g; Polysciences Inc.) was then slowly added, and the reaction was allowed to continue overnight in the dark. The resulting product was precipitated in chilled acetone, rehydrated in DI H_2_O (100 mL), and dialyzed against DI H_2_O at 4 °C for 3 days using dialysis tubing with a molecular weight cutoff (MWCO) of 3.5 kDa (Spectrum Laboratories Inc.), with water changed twice daily. The dialyzed OMA solution was treated with activated charcoal (0.5 mg/100 mL; Neta Scientifics), filtered through a 0.22 μm membrane, frozen at −80 °C overnight, and lyophilized for 10 days until fully dried. ^1^H NMR characterization of O1M20A and O5M20A is shown in **Figure S14** and **Figure S15**, respectively. The actual methacrylation degree determined by ^1^H NMR analysis was 2.8% for O1M20A and 3.2% for O5M20A, respectively, as calculated according to previous reports^[15]^.

### Synthesis of GelMA

GelMA was synthesized according to previously reported protocols with slight modifications^[29]^. Briefly, gelatin type A (10 g; Sigma-Aldrich) was dissolved in PBS (100 mL, pH 7.4) at 50 °C under stirring until fully dissolved. Methacrylic anhydride (10 mL; Sigma-Aldrich) was then added dropwise into the gelatin solution and allowed to react for 1 h at 50 °C. The reaction mixture was subsequently cooled naturally to RT and maintained under stirring overnight. The crude GelMA product was precipitated in chilled acetone, rehydrated in DI H_2_O (100 mL), and dialyzed against DI H_2_O at 50 °C for 7 days using dialysis tubing with an MWCO of 12–14 kDa (Spectrum Laboratories Inc.), with water changed twice daily. The purified GelMA solution was sterilized through a 0.22 μm membrane filter and lyophilized for 10 days. ^1^H NMR characterization of GelMA is shown in **Figure S16**. The actual methacrylation degree was determined to be 54% by ^1^H NMR analysis according to previously reported methods^[30]^.

### Synthesis of GMSs

GMSs were synthesized according to a previously reported protocol with slight modifications^[10c]^. Briefly, gelatin type A (10 g; Sigma-Aldrich) was dissolved in DI H_2_O (90 mL) at 50 °C. The gelatin solution was then added dropwise into preheated olive oil (500 mL, 45 °C) under vigorous stirring at a rate of 10 mL min^-1^ to form a water-in-oil (W/O) emulsion. After stirring for 10 min, the emulsion temperature was gradually reduced to 15 °C while maintaining continuous stirring. After an additional 30 min, chilled acetone (200 mL) was added to the emulsion, and the mixture was stirred for 5 min. The resulting GMSs were collected by filtration and washed five times with acetone to remove residual olive oil. The GMSs were subsequently dried overnight in a fume hood, sterilized under UV irradiation (2 h), and stored at 4 °C until use.

### Synthesis of O1M20A/GMS, O5M20A/GelMA, and O5M20A/GelMA/cells Bioinks

Jammed OMA microgels were prepared according to previously reported protocols with slight modifications^[16a]^. Briefly, O1M20A or O5M20A polymer (2 g) was dissolved in DI H_2_O (100 mL) and subsequently added to a 0.2 M calcium chloride (CaCl_2_) bath (1 L) under vigorous stirring for ionic crosslinking at RT for 4 h, forming OMA hydrogel beads. The hydrogel beads were collected, suspended in 70% ethanol (EtOH; 100 mL), and mechanically fragmented using a blender (Osterizer MFG) operated in pulse mode for 4 min to generate microgels. The fragmented microgel suspension was collected by centrifugation and stored in 70% EtOH at −20 °C until use. On the day of printing, the OMA microgels were washed three times with DI H_2_O containing 0.05% w/v photoinitiator (2-hydroxy-4′-(2-hydroxyethoxy)-2-methylpropiophenone; Sigma-Aldrich), followed by two washes for O1M20A microgels or one wash for O5M20A microgels using low-glucose Dulbecco’s modified Eagle medium (DMEM-LG; Sigma-Aldrich) containing 0.05% w/v photoinitiator, resulting in jammed OMA microgels suitable for extrusion printing. To prepare O1M20A/GMS bioinks, GMSs (40 mg) were mixed with jammed O1M20A microgels (1 mL). To prepare O5M20A/GelMA bioinks, lyophilized GelMA polymer (10 mg) was added to jammed O5M20A microgels (1 mL) and vortexed until fully dissolved. For preparation of O5M20A/GelMA/cell bioinks, freshly harvested cells (NIH3T3, C2C12, hMSCs, or HUVECs; 200 × 10^6^ cells) were mixed with 1 mL of O5M20A/GelMA bioink immediately prior to printing. All bioinks were used immediately after preparation.

### Cell Expansion

hMSCs were isolated according to previously established protocols from bone marrow aspirates^[31]^, which were obtained under a protocol approved by the University Hospitals of Cleveland Institutional Review Board (IRB) and provided in a de-identified form. NIH3T3 fibroblasts (ATCC), C2C12 myoblasts (ATCC), and passage 3 (P3) hMSCs for non-differentiation studies were cultured in GM consisting of DMEM-LG (Sigma-Aldrich) supplemented with 10% fetal bovine serum (FBS; Sigma-Aldrich) and 1% penicillin/streptomycin (P/S; Gibco). Passage 5 (P5) HUVECs (ATCC) were cultured in endothelial growth medium-2 (EGM-2; Lonza). All cells were expanded in T175 tissue culture flasks (Corning) and maintained at 37 °C in a humidified incubator containing 5% CO_2_ until reaching 80–90% confluency. Culture medium was refreshed every other day. On the day of printing, cells were harvested and used for preparation of O5M20A/GelMA/cell bioinks as described above. Passage 4 (P4) hMSCs and passage 6 (P6) HUVECs were used for bioink preparation.

### Rheological Testing

Rheological properties of the bioinks and photocrosslinked hydrogels, including storage (G′) and loss (G″) moduli, shear-thinning behavior, and self-healing behavior, were characterized using a Kinexus Ultra+ rheometer (Malvern Panalytical). Oscillatory measurements were performed using an 8 mm parallel-plate geometry with a 1 mm gap. Bioink or hydrogel samples were loaded between the plates, and all measurements were conducted at 25 °C. Oscillatory frequency sweep tests (0.1–100 Hz at 1% strain) were performed to determine G′ and G″. Shear strain ramp tests (0.01–100% strain at 1 Hz) and shear rate ramp tests (0.1–100 s ¹) were conducted to evaluate shear-thinning behavior. To assess self-healing properties, cyclic strain tests alternating between 1% and 100% strain were performed while monitoring changes in moduli and viscosity. Each strain condition was maintained for 1 min at 1 Hz during cyclic testing. For clarity and improved graph readability, the first three data points immediately following each transition between 1% and 100% strain were omitted from the plotted data. This data presentation does not affect the interpretation of the results or the scientific conclusions.

### Extrusion Printing, Preparation of CytoD and Dead Control Samples, and Construct Culture

Bioinks were loaded into 3 mL syringes fitted with 22-gauge stainless-steel needles (inner diameter: 0.413 mm; McMaster-Carr). Extrusion printing was performed using a BioX bioprinter (CELLINK). Printing parameters for O1M20A/GMS bioinks included a printing speed of 4 mm s^-1^, an extrusion rate of 2.0 μL s^-1^, and an infill density of 40%. For O5M20A/GelMA/cell bioinks, a printing speed of 4 mm s^-1^ and an extrusion rate of 1.5 μL s^-1^ were used. Embedded cell strands were printed as single continuous paths without designated infill density. The base hydrogel construct was printed with a thickness of 1.0 mm, while the embedded cell strand was printed by positioning the needle at a height of 0.5 mm within the base construct. Following printing, constructs were photocrosslinked under UV irradiation using an EXFO Omnicure® S1000 light source (Lumen Dynamics Group) at an intensity of 20 mW cm^-2^ (320–500 nm) for 10–30 s to generate stable hydrogel constructs for subsequent experiments.

To prepare CytoD control samples, CytoD (Invitrogen) was dissolved in sterile dimethyl sulfoxide (DMSO; Sigma-Aldrich) at a stock concentration of 5 mM and freshly supplemented into the culture medium during each medium change to achieve a final concentration of 5 μM. To prepare Dead control samples, harvested cells were suspended in PBS and incubated at 37 °C for 4 h to induce cell death prior to bioink formulation. The resulting dead cells were subsequently used for bioink preparation, extrusion printing, and photocrosslinking to fabricate control constructs.

Strip-shaped constructs were cultured in 12-well plates containing 2 mL of either GM (for NIH3T3-, C2C12-, and hMSC-laden constructs) or EGM-2 (for HUVEC-laden constructs). Larger constructs, including cross-shaped and sheet-shaped constructs, were cultured in 6-well plates containing 6 mL of GM. Half of the culture medium was replaced daily. Constructs were imaged at predetermined time points and analyzed for bending angle quantification according to previously reported methods^[8d, 32]^.

### Degradation and Swelling

Photocrosslinked hydrogels prepared as described above were used for degradation and swelling studies. For degradation testing, hydrogel samples were frozen at −80 °C for 4 h and subsequently lyophilized for 2 days. The dry mass of each sample was recorded as the initial weight (W_i_). The dried hydrogels were then rehydrated and cultured in 5 mL of DMEM-LG at 37 °C. Culture medium was refreshed every other day for O1M20A/GMS hydrogels. In contrast, due to the rapid degradation of O5M20A/GelMA hydrogels, the medium was refreshed every 8 h. At predetermined time points, hydrogels were collected, lyophilized, and weighed to obtain the remaining dry mass (W_d_). Degradation (%) was quantified as mass loss using the following equation: (W_i_-W_d_)/W_i_ × 100%. For swelling studies, hydrogels were cultured in 5 mL of DMEM-LG at 37 °C for 4 h. The diameter of the swollen hydrogel (D_s_) was measured and compared with the original hydrogel diameter (D_o_). Swelling was calculated using the following equation: D_s_/D_o_.

### Chondrogenesis and Osteogenesis Studies

P3 hMSCs were expanded for both chondrogenesis and osteogenesis studies. hMSCs prescreened for high chondrogenic differentiation potential from donor 1 were used for chondrogenesis studies, whereas hMSCs prescreened for high osteogenic differentiation potential from donor 2 were used for osteogenesis studies. For chondrogenic expansion, hMSCs were cultured in DMEM-LG supplemented with prescreened 10% FBS, 1% P/S, and 10 ng mL^-1^ fibroblast growth factor-2 (FGF-2; R&D Systems). For osteogenic expansion, hMSCs were cultured in high-glucose DMEM (DMEM-HG) supplemented with 10% FBS and 1% P/S. Cells were harvested at approximately 80% confluency and used for bioink preparation.

After printing using P4 hMSCs, constructs for chondrogenesis were cultured in chondrogenic differentiation medium consisting of DMEM-LG supplemented with 10% insulin-transferrin-selenium plus (ITS^+^; Fisher Scientific), 1% non-essential amino acids (NEAAs; Gibco), 1% P/S, 100 mM sodium pyruvate (Fisher Scientific), 100 nM dexamethasone (Sigma-Aldrich), L-ascorbic acid phosphate (Wako USA), and 10 ng mL^-1^ transforming growth factor-β1 (TGF-β1; PeproTech). Constructs for osteogenesis were cultured in osteogenic differentiation medium consisting of DMEM-HG supplemented with 10% FBS, 1% P/S, 10 mM β-glycerophosphate (Sigma-Aldrich), 50 μM ascorbic acid (Sigma-Aldrich), 100 nM dexamethasone (Sigma-Aldrich), and 100 ng mL^-1^ bone morphogenetic protein-2 (BMP-2; GeneScript). Constructs were collected for biochemical, mechanical, and histological analyses at D21 for chondrogenesis and D28 for osteogenesis.

For biochemical analyses, chondrogenic constructs at D21 and osteogenic constructs at D28 were collected and homogenized. Chondrogenic constructs were immersed in 1 mL papain digestion solution (pH 6.5) containing 50 μg mL^-1^ papain (Sigma-Aldrich), 2 mM L-cysteine (Sigma-Aldrich), 50 mM sodium phosphate (Sigma-Aldrich), and 2 mM EDTA (Fisher Scientific). Osteogenic constructs were immersed in 1 mL CelLytic^TM^ buffer (Sigma-Aldrich). Samples were homogenized on ice at 35,000 rpm for 1 min using a TH homogenizer (Omni International). Chondrogenic samples were subsequently digested overnight at 65 °C and centrifuged at 500× g to collect supernatants for DNA and GAG analysis. Osteogenic samples were centrifuged at 500× g to collect supernatants for DNA and ALP analysis, while the remaining pellets were treated with 1.2 N hydrochloric acid (HCl) overnight at 4 °C for calcium quantification^[33]^.

DNA, GAG, ALP, and calcium analyses were performed according to previously established methods^[31a, 32]^. DNA content was quantified using a PicoGreen dsDNA fluorescence assay (Invitrogen) with excitation at 480 nm and emission at 520 nm using a SpectraMax iD5 plate reader (Molecular Devices). GAG content was measured using a 1,9-dimethylmethylene blue (DMMB; Sigma-Aldrich) assay at 595 nm. ALP activity was determined by measuring absorbance of p-nitrophenyl phosphate (pNPP)-treated samples after incubation at 37 °C for 30 min at 405 nm. Calcium content was quantified using a calcium assay kit (Pointe Scientific) with absorbance measured at 570 nm.

### MTJ Regeneration

To mimic MTJ architecture, NIH3T3 fibroblasts and C2C12 myoblasts were used. NIH3T3 fibroblasts were incorporated into O1M20A/GMS bioinks at a density of 10 × 10^6^ cells mL^-1^ bioink and printed as the base hydrogel construct. Subsequently, O5M20A/GelMA/cell bioinks containing C2C12 cells (200 × 10^6^ cells mL^-1^ bioink) were embedded and printed within the base construct as described above. Base hydrogel controls consisted of NIH3T3-laden hydrogel constructs without embedded cell strands. All constructs were cultured in MTJ differentiation medium consisting of GM supplemented with 2% horse serum (Gibco) and 1% ITS^+[34]^.

For fluorescence tracking, NIH3T3 cells and C2C12 cells were labeled using CellTracker Green CMFDA and CellTracker Red CMTPX dyes (Fisher Scientific), respectively. Stock solutions (10 mM) were prepared in sterile DMSO and diluted in serum-free DMEM-LG to obtain staining media containing 20 μM CellTracker Green or 10 μM CellTracker Red. Harvested cells were incubated in 20 mL staining medium in T175 flasks at 37 °C for 30 min (CellTracker Green) or 15 min (CellTracker Red). Cells were subsequently collected by centrifugation and used for bioink preparation, printing, and culture. Samples collected at predetermined time points were imaged using a TE300 fluorescence microscope (Nikon) equipped with an AmScope MU1403 camera (AmScope).

### Live/Dead Staining and Histological Analysis

Constructs collected at predetermined time points were stained with fluorescein diacetate (FDA; Sigma-Aldrich) and ethidium bromide (EtBr; Sigma-Aldrich) to visualize live (green) and dead (red) cells, respectively. Constructs were incubated in staining solution in DMEM-LG at RT for 3 min prior to fluorescence imaging.

Histological analyses were performed according to previously established protocols^[10c, 10d]^. Constructs were fixed in 10% neutral buffered formalin (NBF) at 4 °C overnight, dehydrated, infiltrated with paraffin (Epredia), embedded into paraffin blocks, and sectioned into 7 μm slices using an RM2255 microtome (Leica Biosystems). For H&E staining, sections were stained with Mayer’s hematoxylin (Fisher Scientific) for 2 min followed by eosin-phloxine solution (Electron Microscopy services) for 30 s. For F-actin staining, sections were permeabilized with Triton X-100 (0.5% w/v; Fisher Scientific), blocked with bovine serum albumin (BSA; 1% w/v; Fisher Scientific), stained with Alexa Fluor 488-conjugated phalloidin (Fisher Scientific) for 1 h in the dark, and counterstained with DAPI (Sigma-Aldrich) for 5 min. For E-cadherin staining, sections were permeabilized with Triton X-100 (0.5% w/v), blocked with donkey serum (1% w/v; Jackson ImmunoResearch), incubated with primary mouse anti-E-cadherin antibody (Fisher Scientific) overnight at 4 °C, followed by Alexa Fluor 488-conjugated secondary antibody staining and DAPI counterstaining. SafO/FG, Alcian blue (pH 1.0), TBO, and ARS staining were performed using reported protocols^[10c, 10d, 35]^ to evaluate cartilage-like and bone-like matrix deposition. Stained sections were imaged using the Nikon fluorescence microscope.

### Statistics

All quantitative data are presented as mean ± standard deviation (SD) from triplicate samples (*N* = 3). Statistical analyses were performed using one-way analysis of variance (ANOVA) followed by Tukey’s honestly significant difference (HSD) post hoc test for multiple comparisons. Differences were considered statistically significant at *p* < 0.05 unless otherwise specified.

## Acknowledgements

The authors gratefully acknowledge funding support from the U.S. Department of Veterans Affairs, Veterans Health Administration, Office of Research and Development, Rehabilitation Research and Development Service under award number RX004288, and from the National Institutes of Health National Institute of Arthritis and Musculoskeletal and Skin Diseases under award number R01AR081448. A.F.C. acknowledges FCT for doctoral grant 2021.04817.BD and FLAD for the mobility grant (Project No. 2025/0023). A.F.C, M.B.O., and J.F.M. acknowledge the project CICECO-Aveiro Institute of Materials, UID/50011/2025 (DOI 10.54499/UID/50011/2025) & LA/P/0006/2020 (DOI 10.54499/LA/P/0006/2020) & UID/PRR/50011/2025 (DOI 10.54499/UID/PRR/50011/2025), financed by national funds through the FCT/MCTES (PIDDAC). The contents of this publication are solely the responsibility of the authors and do not necessarily represent the official views of the Department of Veterans Affairs or the National Institutes of Health. The authors also thank Dr. Kaelyn L. Gasvoda of the Mayo Clinic and Dr. Fang Tang of Northwestern University for their assistance in creating the schematic illustrations of the hydrogel constructs and morphogenetic architectures presented in this study.

## Data Availability Statement

The data that support the findings of this study are available from the corresponding author upon reasonable request.

## Movie Captions

**Movie S1.** A NIH3T3 strand-laden hydrogel strip (10 s UV) at D7 exhibited softness while maintaining sufficient mechanical robustness to withstand vigorous spatula stirring in PBS (4x speed).

**Movie S2.** A GM-cultured hMSC strand-laden hydrogel strip at D21 exhibited sufficient mechanical robustness to withstand vigorous spatula stirring in PBS (2x speed).

**Movie S3.** A CPM-cultured hMSC strand-laden hydrogel strip at D21 exhibited sufficient mechanical robustness to withstand vigorous spatula stirring in PBS (1x speed).

**Movie S4.** A GM-cultured hMSC strand-laden hydrogel strip at D21 could not withstand compression with tweezers (2.5x speed).

**Movie S5.** A CPM-cultured hMSC strand-laden hydrogel strip at D21 exhibited high elasticity and complete shape recovery after full compression with tweezers (3x speed).

**Movie S6.** A trachea-mimicking tubular cartilage construct in PBS being manipulated with a spatula. Note that one ring detached from the construct during transfer from CPM to PBS (3x speed).

**Movie S7.** A GM-cultured hMSC strand-laden hydrogel strip at D28 exhibited sufficient mechanical robustness to withstand vigorous spatula stirring in PBS (2x speed).

**Movie S8.** An OM-cultured hMSC strand-laden hydrogel strip at D28 exhibited sufficient mechanical robustness to withstand vigorous spatula stirring in PBS (3x speed).

**Movie S9.** An OM-cultured hMSC strand-laden hydrogel strip at D28 exhibited complete shape recovery after full compression with tweezers (3x speed).

**Movie S10.** A tubular bone construct in PBS being manipulated with a spatula (3x speed).

