## Supporting Information for "Self-Actuating 4D Cell-Strand Bioprinting"

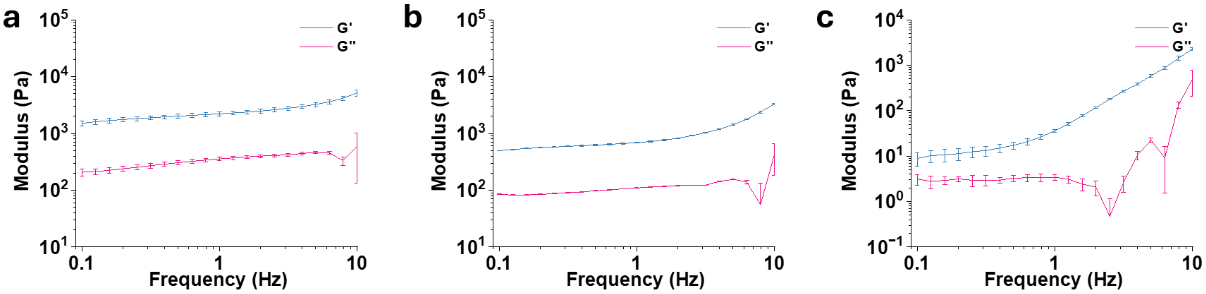


**Figure S1. Rheological characterization of bioinks.** Storage modulus (G′) and loss modulus (G″) of (a) O1M20A/GMS bioinks, (b) O5M20A/GelMA bioinks, and (c) O5M20A/GelMA/cells bioinks as functions of oscillatory frequency. NIH3T3 cells were encapsulated at a density of 200 million cells $mL^{-1}$ bioink. Data are presented as mean ± standard deviation (±SD), $N = 3$.

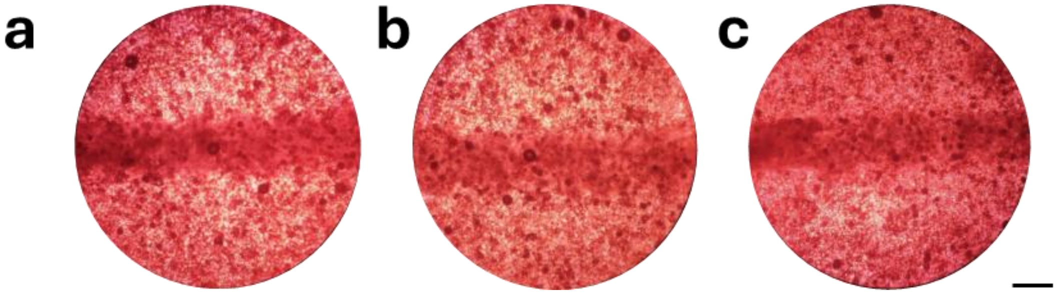


**Figure S2.** Representative images of printed cell strands within the O1M20A/GMS base hydrogel containing (a) NIH3T3 cells, (b) C2C12 cells, and (c) hMSCs. Scale bar = 0.5 mm.

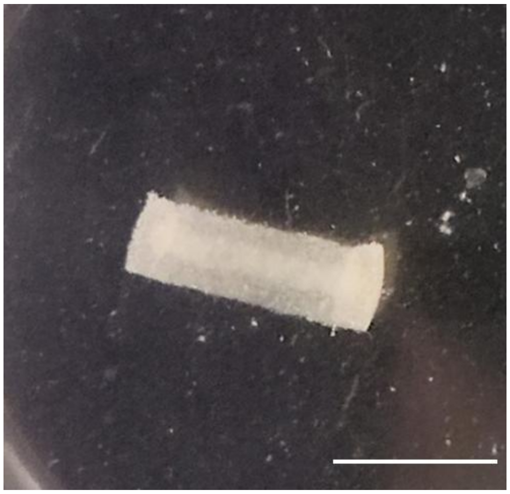

**Figure S3.** The cell strand remained intact and clearly visible within a hydrogel strip (20 s UV, NIH3T3) after 7 days of culture in GM. Scale bar = 5 mm.

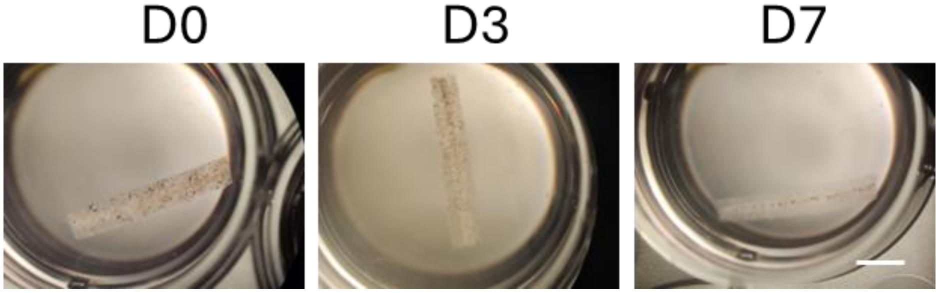


**Figure S4.** Shape transformation of HUVEC-laden hydrogel strips during culture in endothelial growth medium. Scale bar = 5 mm.

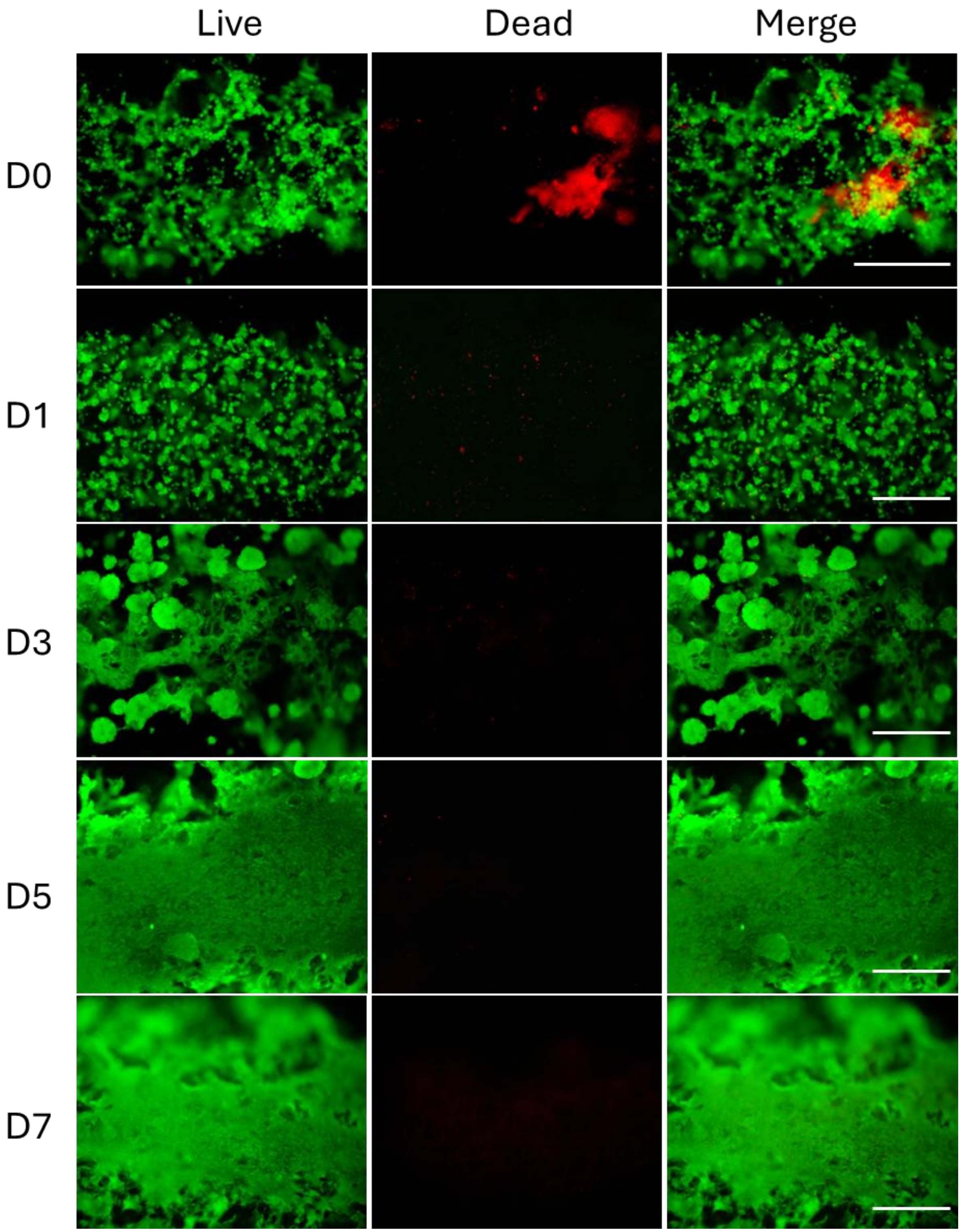


**Figure S5.** Magnified live/dead staining images of NIH3T3 strand-laden hydrogel strips at different times. Scale bars = 0.25 mm.

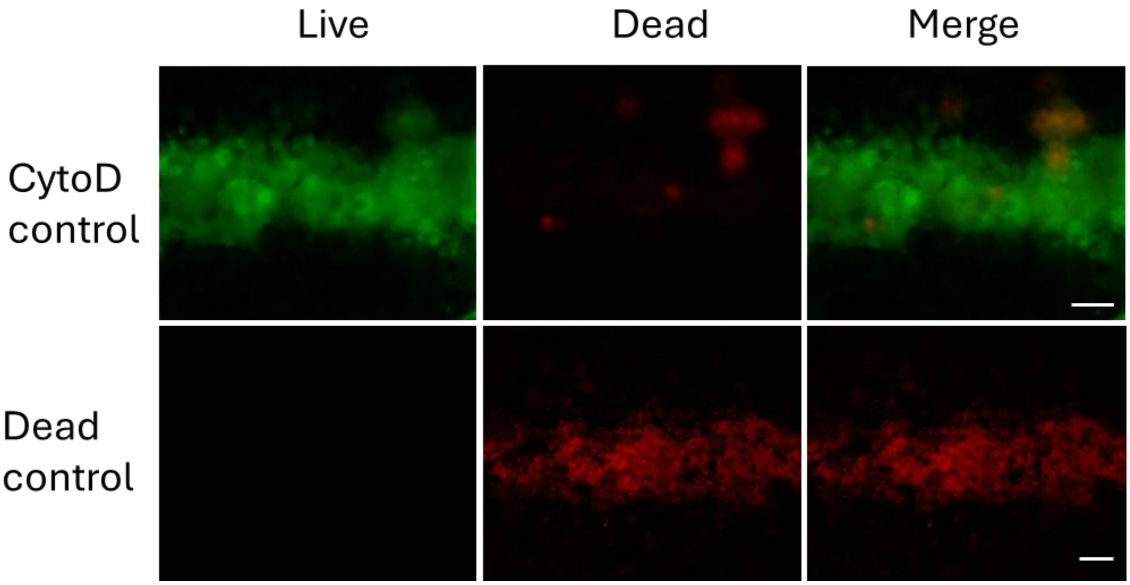


**Figure S6.** Representative live/dead staining images of CytoD and Dead control strips at D0. Scale bars = 0.25 mm.

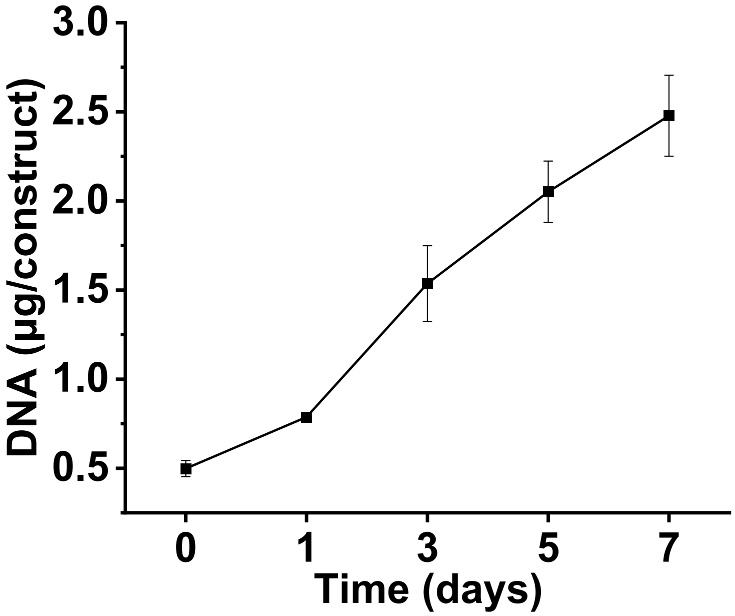


**Figure S7.** Changes in DNA content of NIH3T3 strand-laden hydrogel strips during culture in GM. Data are presented as mean ± standard deviation (±SD), $N = 3$.

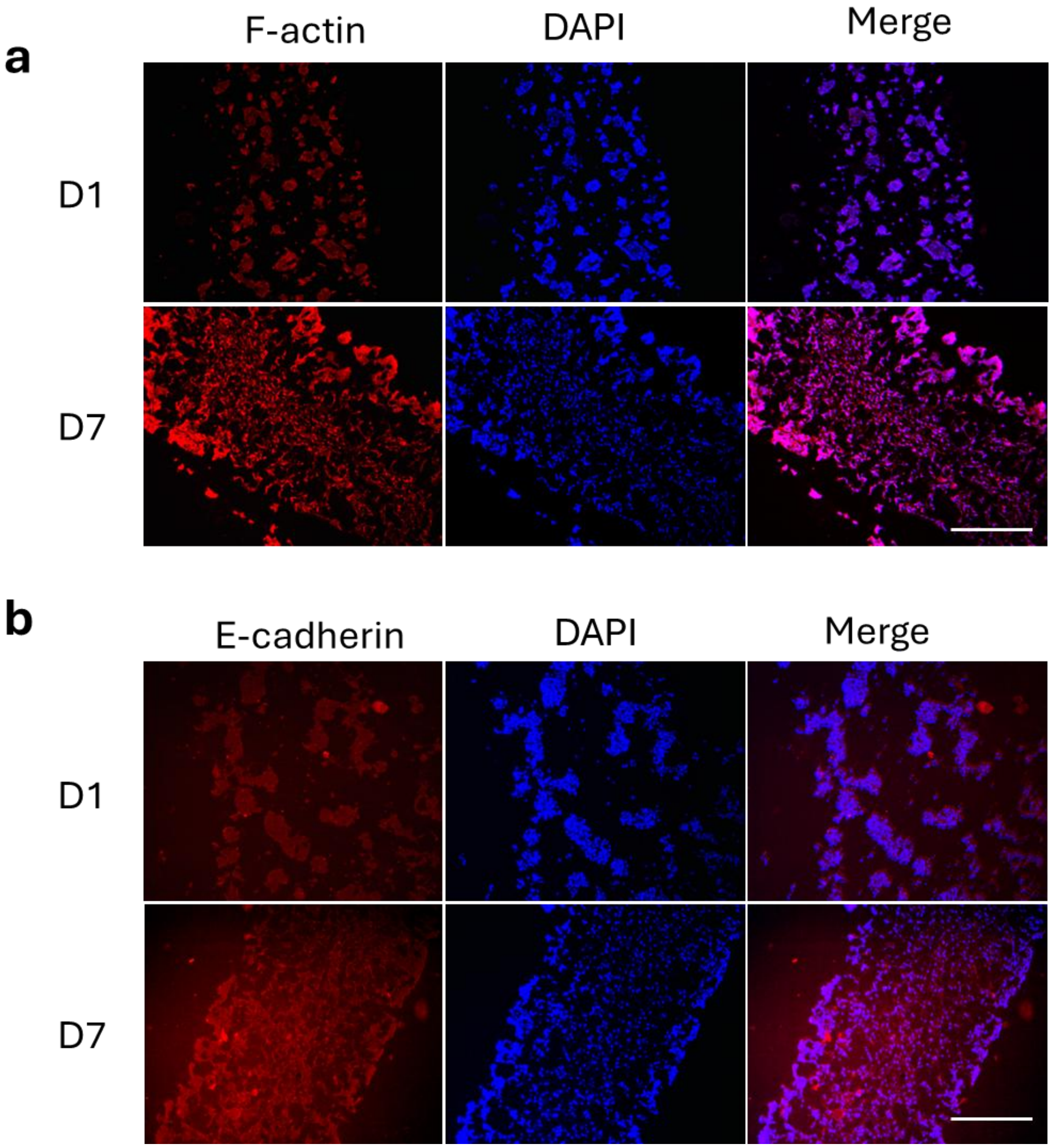


**Figure S8. Histological staining of NIH3T3 construct-laden constructs at D1 and D7**. Photomicrographs of (a) F-actin and (b) E-cadherin stained sections. Scale bars = 0.25 mm.

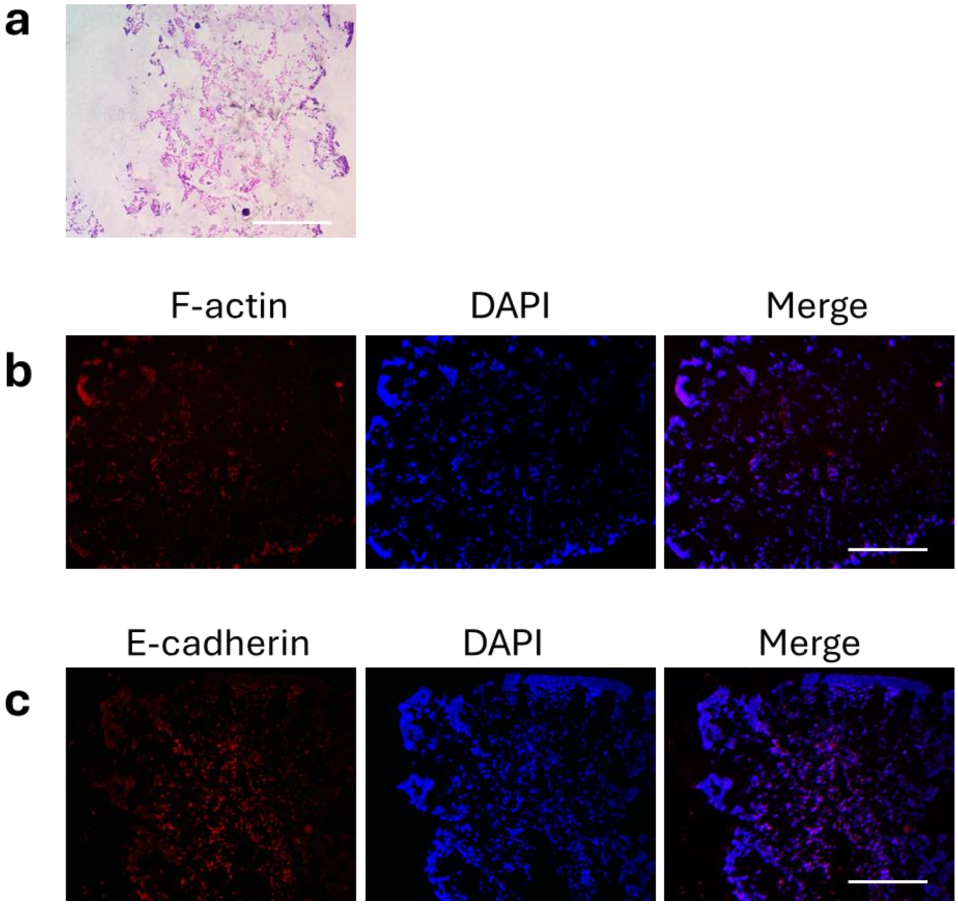


**Figure S9. Histological characterization of CytoD control strips at D7.** Photomicrographs of (a) H&E, (b) F-actin, and (c) E-cadherin stained sections. Scale bars = 0.25 mm.

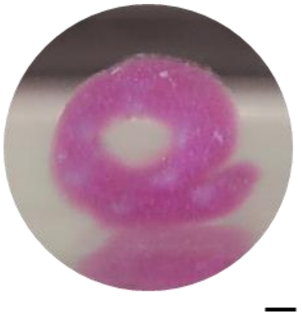

**Figure S10.** Front view of the 7-strand tubular construct at D7 of culture GM, imaged in PBS. Scale bar = 1 mm.

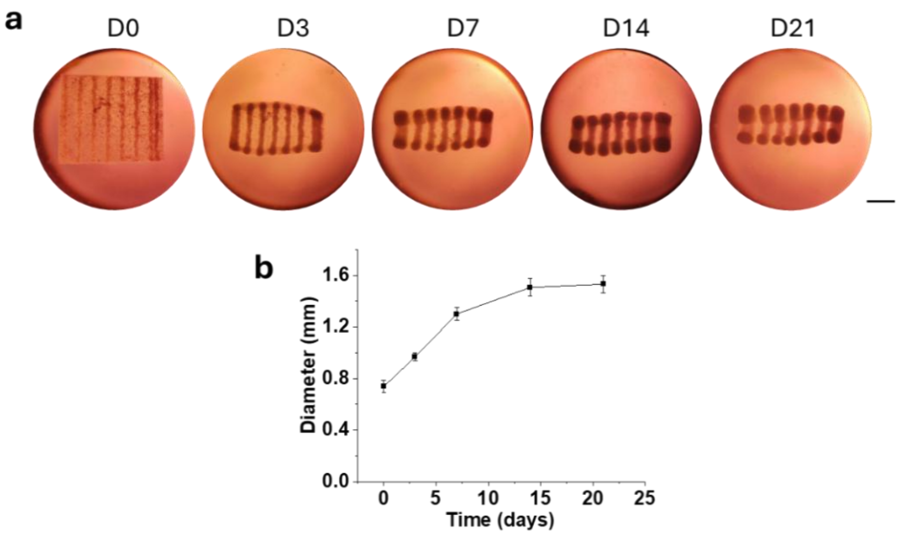


**Figure S11. Characterization of 7-strand hMSC-laden hydrogel constructs during 21 days of culture in CPM.** (a) Representative images showing progressive shape morphing over culture time. (b) Changes in strand diameter over time. Scale bar = 5 mm. Data are presented as mean ± standard deviation (±SD), $N = 3$.

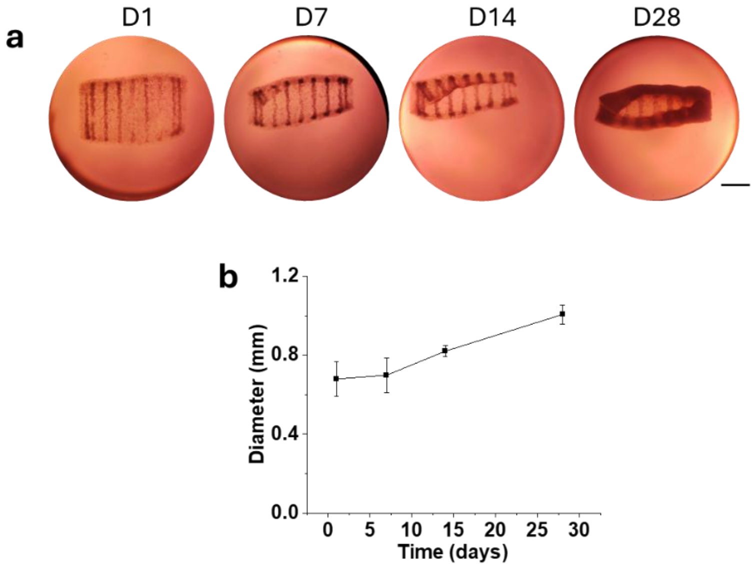


**Figure S12. Characterization of 7-strand hMSC-laden hydrogel constructs during 28 days of culture in OM.** (a) Representative images showing progressive shape morphing over culture time. (b) Changes in strand diameter over time. Scale bar = 5 mm. Data are presented as mean ± standard deviation (±SD), $N = 3$.

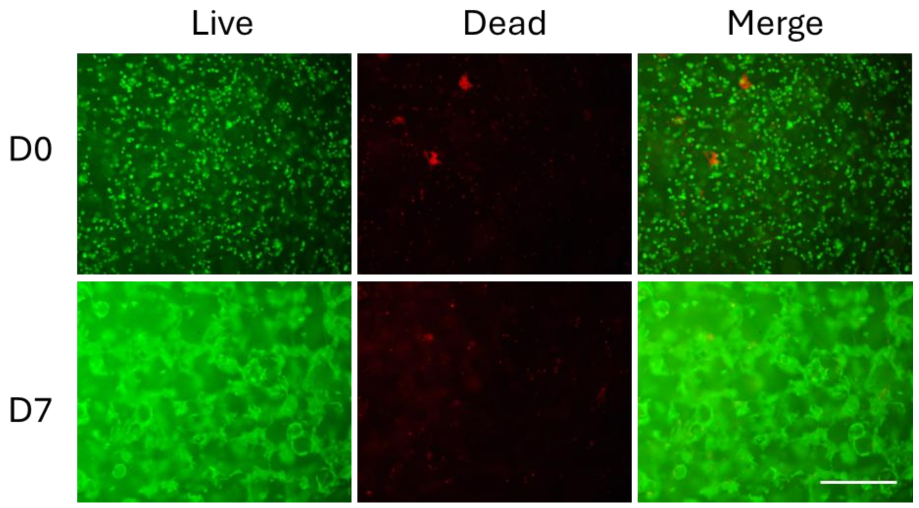


**Figure S13.** Live/dead staining of NIH3T3 cells encapsulated within the base hydrogel. Scale bar = 0.25 mm.

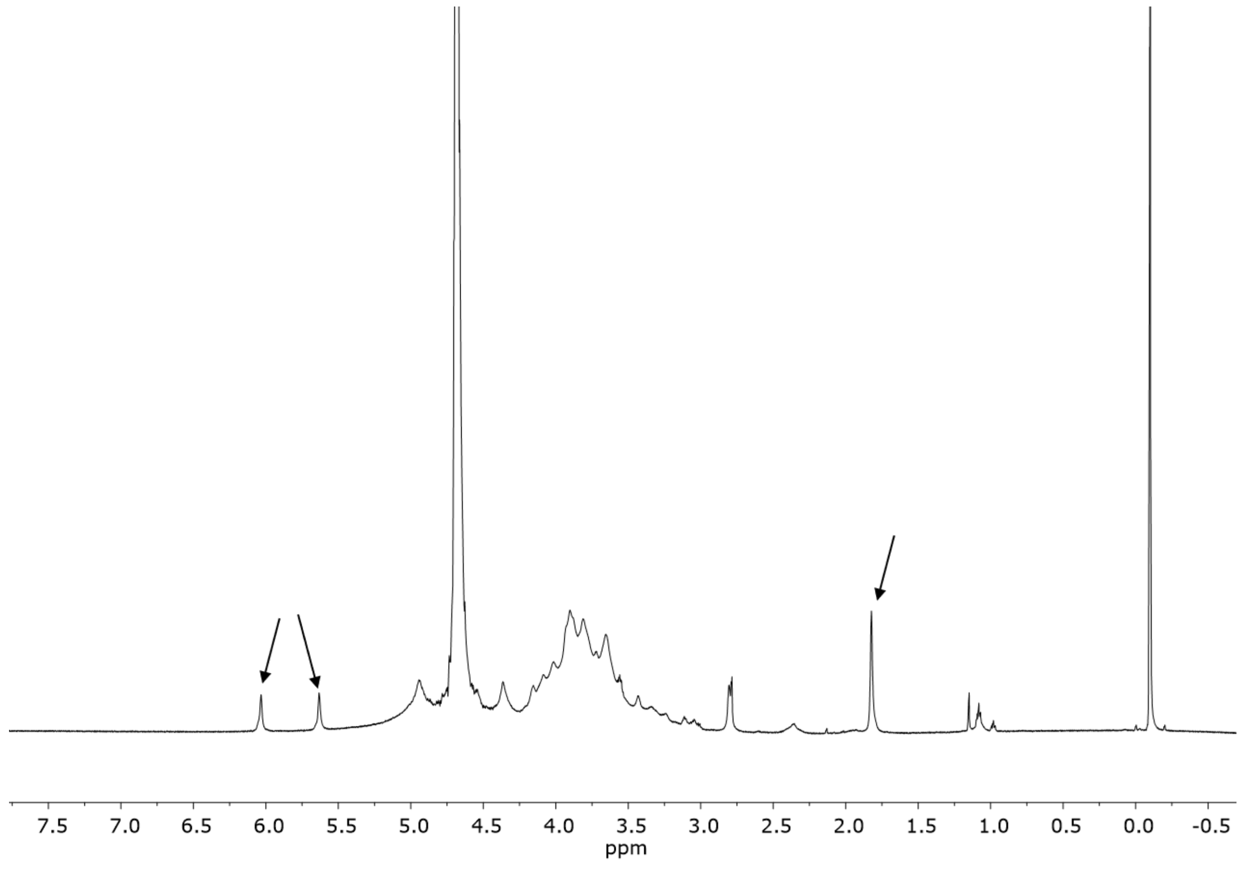


**Figure S14.** $^1H$ NMR spectrum of O1M20A in $D_2O$. Arrows indicate the characteristic methacrylate proton peaks.

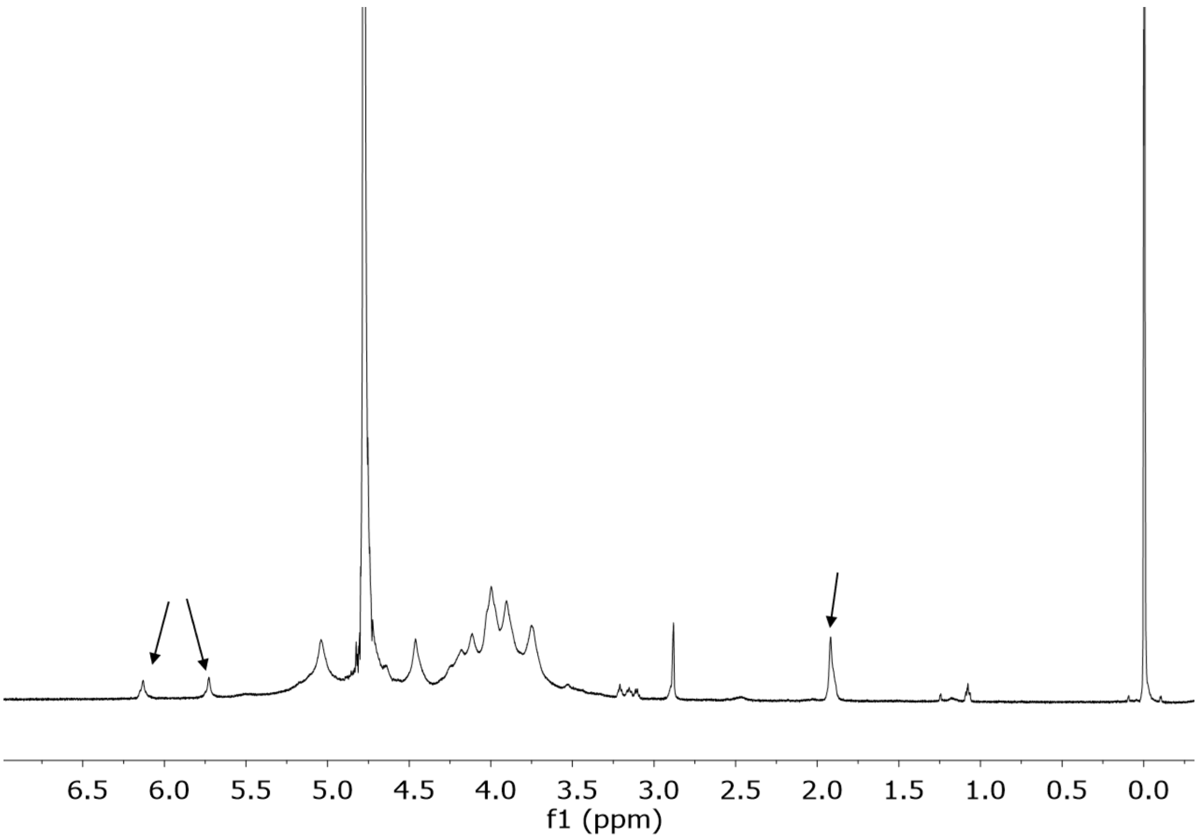


**Figure S15.** [1]H NMR spectrum of O5M20A in $D_2O$. Arrows indicate the characteristic methacrylate proton peaks.

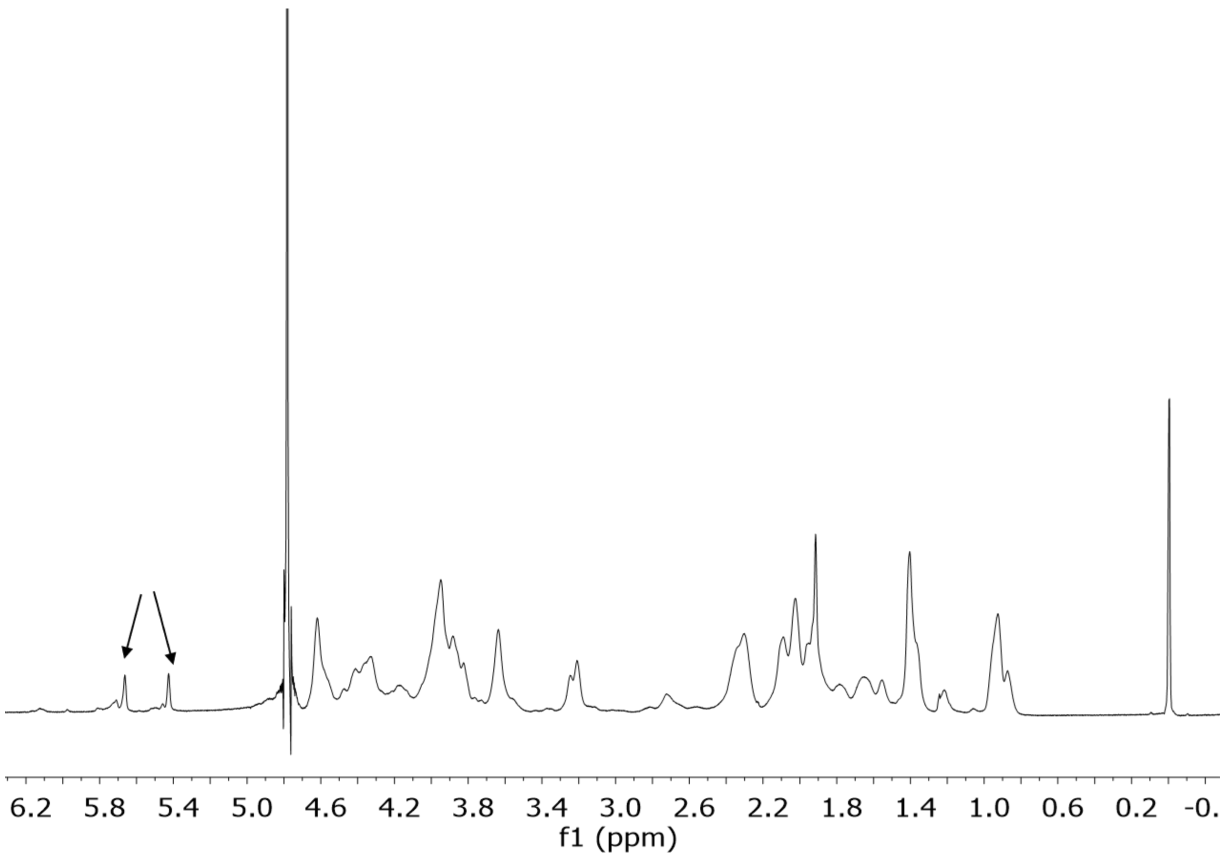


**Figure S16.** $^{1}H$ NMR spectrum of GelMA in $D_2O$. Arrows indicate the characteristic methacrylate proton peaks.
